# Highly effective antifibrotic compound discovered as a contaminant in cefixime preparations: preliminary characterization

**DOI:** 10.1101/2025.08.28.672907

**Authors:** Branko Stefanovic

## Abstract

There are no approved drugs for specific treatment of excessive type I collagen synthesis in organ fibrosis. The discovery that biosynthesis of type I collagen in fibrosis is regulated by binding of LARP6 to the unique sequence found in type I collagen mRNAs, the 5’ stem-loop (5’SL), prompt the search for small molecule inhibitors of LARP6 binding. A contaminant in some commercial preparations of the third-generation cephalosporin, cefixime, has been discovered that inhibits LARP6/5’SL interaction in vitro and type I procollagen secretion by cells and organoids in culture. The activity, termed ATO-OA, forms stable non-colloidal nano-entities (NE). ATO-OA NE inhibit the binding of LARP6 to 5’SL with the IC_50_ of 3-4 µg/ml and are equally effective in the dissociation of preassembled LARP6/5’SL complex as in the inhibition of its formation. ATO-OA NE target the RRM domain of LARP6 to alter its conformation. In cultured cells, ATO-OA NE suppresses type I procollagen secretion into the cellular medium and causes intracellular retention of the protein. The effect of ATO-OA NE *in vivo* is LARP6 dependent, because the LARP6 knockout cells or the cells which make type I procollagen without 5’SL show no effect of ATO-OA. In cultured organoids of human pancreatic adenocarcinoma cells ATO-OA NE diminished type I collagen production at ∼62.5 µg/ml. These results suggest that the discovery of ATO-OA NE as potent and specific inhibitor of LARP6 may be a breakthrough towards development of antifibrotic drugs directly targeting type I collagen biosynthesis.

## Introduction

Excessive biosynthesis of type I collagen is the main contributor to pathogenesis of organ fibrosis [1–3]. The goal of any relevant treatment for organ fibrosis must be to reduce type I collagen deposition in the tissues. However, the current efforts to find antifibrotic drugs are limited to the approaches that indirectly suppress type I collagen biosynthesis by targeting proinflammatory mechanisms or pleiotropic cell signaling pathways, such as TGF-β, Wnt, mTOR, CTGF, PDGF [4–17]. However, it has also been well documented that the binding of RNA binding protein LARP6 to the mRNAs encoding for type I procollagen is a critical regulatory step that accelerates type I collagen biosynthesis in fibrosis [18–30]. Mice in which LARP6 dependent type I collagen biosynthesis is obliterated, are resistant to development of hepatic fibrosis and had reduced arterial stiffness in a vascular injury model [27, 31]. These findings indicated that inhibiting LARP6 binding to collagen mRNAs can be an effective strategy to suppress fibrosis progression in vivo [32].

Type I procollagen is composed of two α1 polypeptides and one related α2 polypeptide [33]. The mRNAs encoding these polypeptides (COL1A1 and COL1A2) have in their 5’ UTR a secondary structure, designated as collagen mRNA 5’ stem-loop (5’SL) [34, 35]. The 5’SL has unique sequence and structure and is found exclusively in type I collagen mRNAs and in type III collagen mRNA. The RNA binding protein LARP6 binds 5’SL with high affinity and regulates translation of type I procollagen mRNAs by restricting their random loading with polysomes and converging the translation of type I collagen pro-peptides into distinct structures on the endoplasmic reticulum membrane, named collagenosomes [36–38]. The biosynthesis in collagenosomes accelerates the folding of three type I collagen polypeptides into procollagen [38], and results in its rapid secretion into the extracellular matrix. This mechanism increases the fractional synthesis rate of type I collagen and contributes to organ fibrosis [39].

LARP6 belongs to the LARP superfamily of translational regulators but is the only family member which can recognize collagen 5’SL sequence with high affinity and sequence specificity [34, 35, 40]. LARP6 has two domains that are involved in 5’SL binding, LA and RRM, and recognizes noncanonically base paired nucleotides in the bulge of 5’SL [35]. We have previously described two LARP6 binding inhibitors and characterized one of these inhibitors in animal models of hepatic fibrosis [32, 41]. This manuscript describes the discovery of a novel activity that inhibits LARP6 binding to 5’SL in vitro and type I procollagen biosynthesis in human cells and organoids.

## Results

### Conserved 5’SL of type I collagen mRNAs

Figure 1 compares the sequence and secondary structure of 5’SL of human COL1A1 mRNA and COL1A2 mRNA. The 5’SL includes the start codon in both mRNAs. The nucleotides of 5’SL RNA involved in recognition of LARP6 are shown in red or circled in Fig 1 [34]. These nucleotides form noncanonical base pairs [35] and are identical between A1 and A2 5’SL sequences, which bind LARP6 with similar affinity. To assess LARP6 binding to 5’SL RNA the A1 5’SL RNA and A2 5’SL RNA sequences shown in Fig 1 were used throughout this manuscript.

**Figure 1.**
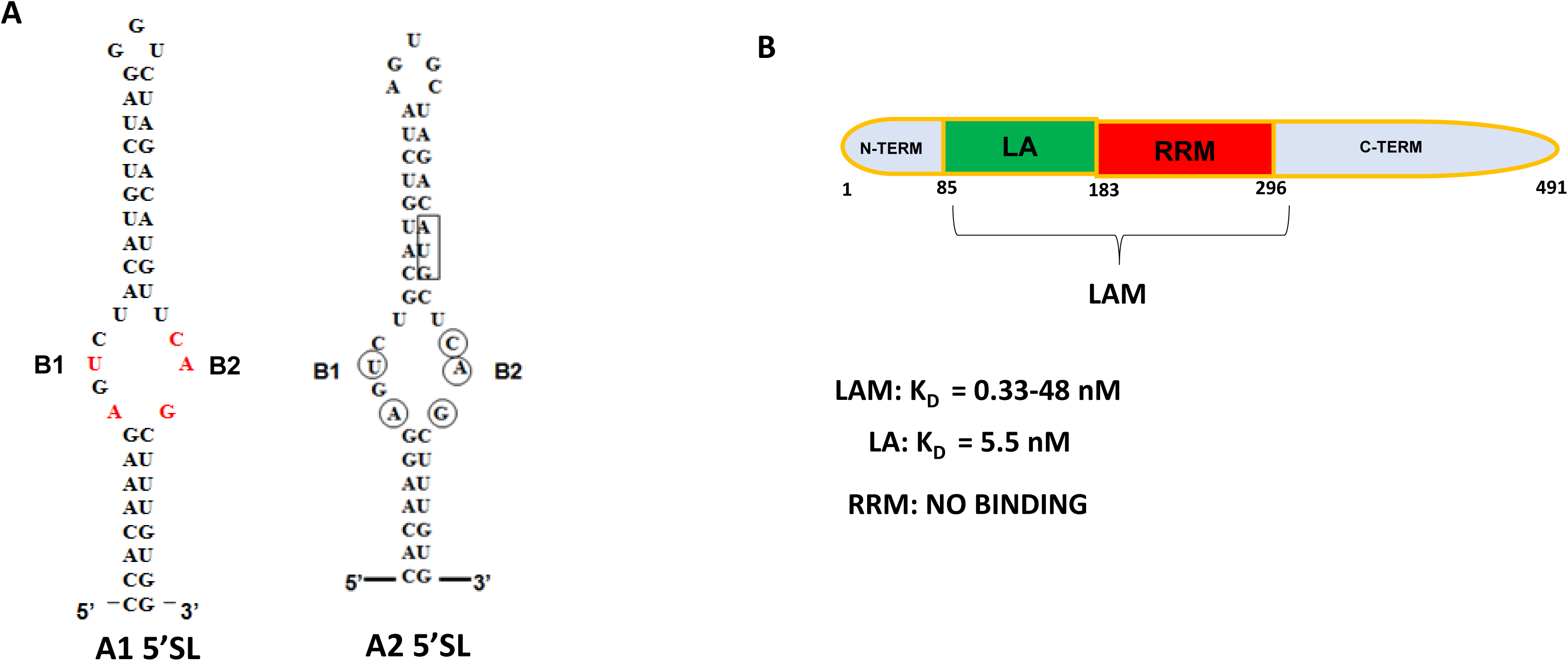
Secondary structures of COL1A1 and COL1A2 5’ stem loops (5’SL) and schematic representation of human LARP6 domains. A. COL1A1 and COL1A2 5’ SL RNAs. Nucleotides involved in LARP6 recognition are shown in red for A1 5’SL and circled for A2 5’SL. B1 and B2; region of noncanonical base pairing. B. Domains of LARP6. LA and RRM in tandem are termed the LA-module (LAM). Affinity of binding to A1 5’ SL of the domain is indicated.

LARP6 has two domains that participate in 5’SL binding; the LA domain (green in Fig 1) and the RRM domain (red in Fig 1), while the rest of the protein is dispensable for binding [34, 40]. When both domains were expressed in tandem they are referred to as the La-module (LAM), while the individual domains are referred to as LA or RRM [42]. The LA domain alone can bind 5’SL, while the RRM cannot, however the presence of RRM increases the affinity of binding. Therefore, LAM binds 5’SL with 5-10 fold greater affinity than LA alone, as indicated in Fig 1 [23, 24].

### An inhibitor of LARP6 binding discovered in commercial preparations of cefixime

During our screening of FDA approved drugs for inhibitors of LARP6 binding to 5’SL RNA we observed that a sample of the third-generation cephalosporin, cefixime, inhibited *in vitro* interaction of LAM and 5’SL RNA at higher concentrations. When several other commercial preparations of cefixime were tested, we found that some preparations contained the inhibitory activity and some did not. Figure 2A shows *in vitro* binding of LAM to A1 5’ SL RNA analyzed by gel mobility shift experiments in the presence of various concentrations of three different cefixime preparations from commercial sources. In the gel mobility shift assays free RNA was resolved as monomer and dimer; dimer of 5’SL RNA spontaneously appears because of the palindromic sequence of 5’SL RNA. LAM/A1 RNA complexes were resolved as 2-3 bands of lower electrophoretic mobility [24, 34]. The decrease in intensity of LAM/RNA complexes accompanied with the increase in amount of free RNA was used as a readout of inhibition.

**Figure 2.**
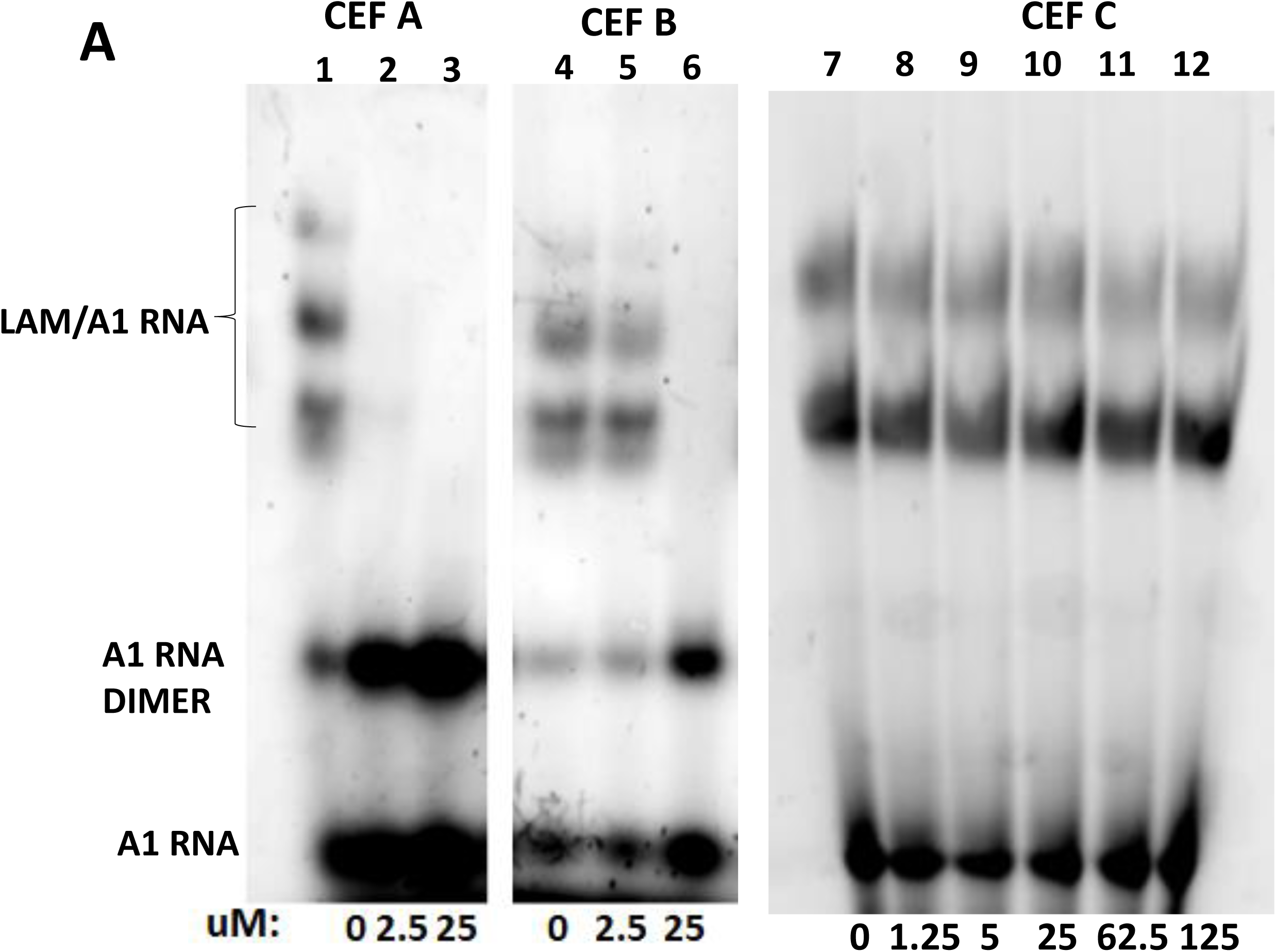
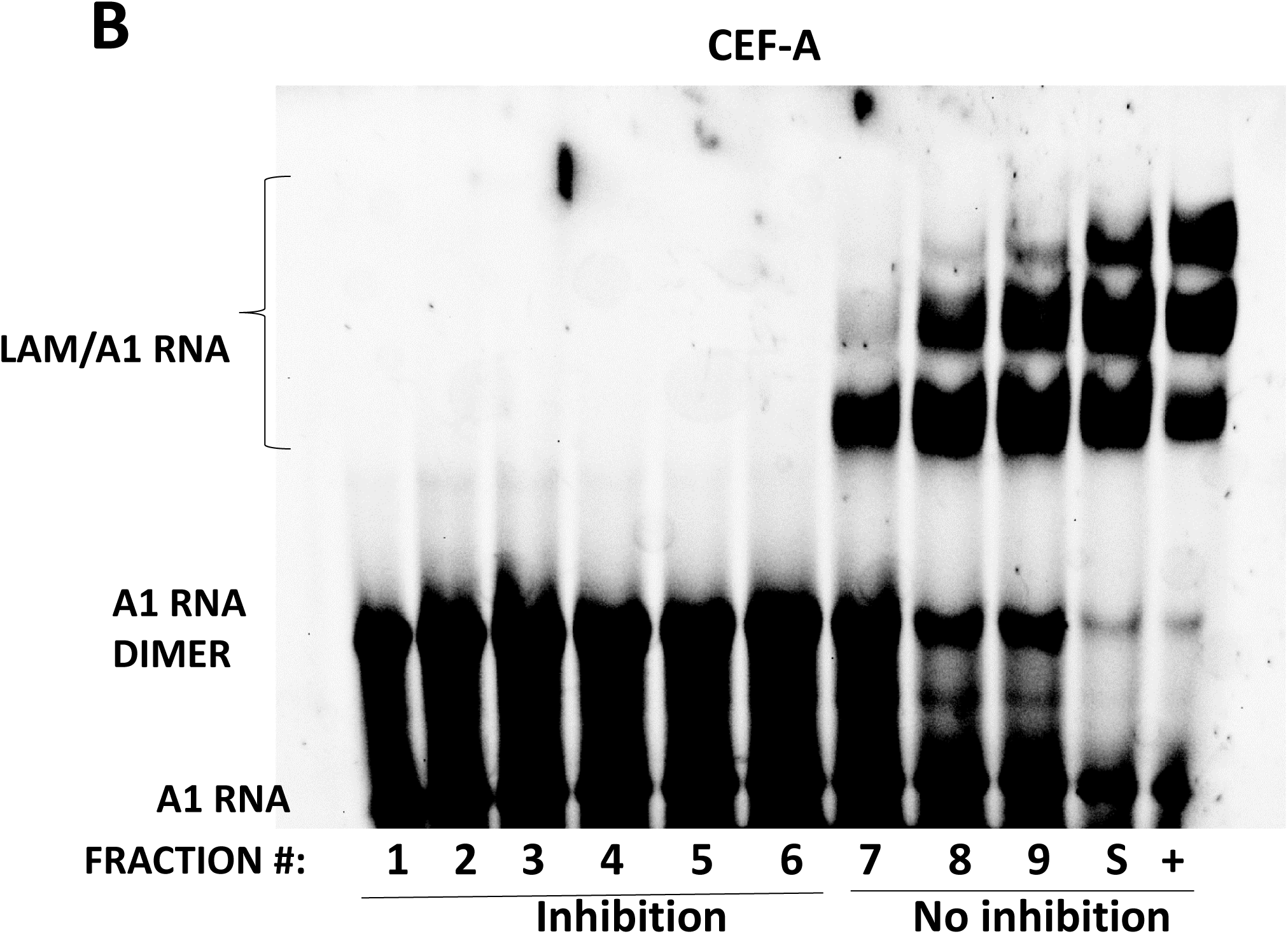
A contaminant present in some commercial preparations of cefixime inhibits LARP6 binding. A. Comparison of inhibitory activity of cefixime from three different commercial sources (CEF A, CEF B, CEF C). Gel mobility shift using recombinant LAM and A1 5’SL RNA and the indicated concentrations of CEF A (lanes 1-3), CEF B (4-6) and CEF C (lanes 7-12). Mobility of free RNA monomer (A1 RNA), free RNA dimer (A1 RNA DIMER) and LAM/5’ SL RNA complex (LAM/A1 RNA) is indicated. B. Fractionation of CEF A by stepwise solvent shift. Fractions collected after each solvent shift were analyzed for inhibition of LAM binding by gel mobility shift. Active fractions (1-6) and inactive fractions (7-9) are indicated. S, supernatant after fraction 9. +, positive control for binding.

Fig 2A shows that the cefixime preparation A inhibited LAM binding at concentration of 2.5 µM and higher (lanes 2 and 3), while cefixime preparation B inhibited at concentration of 25 µM (lane 6) and cefixime preparation C was inactive up to 125 µM (lanes 7-12). This indicated that the inhibitory activity could not be ascribed to cefixime, but to a contaminant present in various amounts in different preparations.

### Separation of the inhibitor from cefixime

To separate the inhibitor from cefixime CEF-A was fractionated by stepwise solvent shift. After each step of adding 10% volume of water to the DMSO solution of CEF-A a precipitate was formed, which was collected and re-dissolved in DMSO or in phosphate buffer pH 6. Nine fractions and the supernatant after the fraction 9 were obtained and equal volume of each fraction was analyzed for inhibition of LAM binding (Fig 2B). Fractions 1-6 showed strong inhibition of binding, fraction 7 was partially active, while fractions 8, 9 and the supernatant after fraction 9 (S) were inactive. Mass spec analysis of the fractions revealed the presence of an ion of [M+Z]^+^ =287.0444 in early fractions, which we termed ATO-OA. The structure and synthesis of ATO-OA will be described in the follow up manuscript. Here, show the results with the ATO-OA purified from cefixime preparations.

### ATO-OA inhibitor forms non-colloidal nano-entities (NE)

The result of dynamic light scattering (DLS) [43] measurement of the ATO-OA hydrodynamic radius in aqueous solution is shown in Fig 3A. Two peaks with the mean hydrodynamic radius of 8.62 nM and 47.3 nM were resolved (dark blue peaks), suggesting that ATO-OA exists as nano-entities (NE) [44]. The 8.62 nM peak contained 90% of mass, indicating that this is the predominant form of ATO-OA NE. The 47.27 nM peak contained only 6.6% of mass. When Triton X-100 was added at concentration of 0.1%, there was a shift of the 8.62 mM peak to 5.0 nM (light blue peak); this peak contained 89% of the mass. With Triton X-100, the 47.27 nM peak shifted to 33.5 nM and contained 6.3% of mass (light blue peak). The persistence of NE with high hydrodynamic radius in the presence of Triton X-100 suggested that ATO-OA NE are not colloidal aggregates [45, 46], because Triton X-100 at 0.1% completely dissolves colloidal aggregates [47]. The ghost peak seen with Triton X-100 had no mass associated with it.

**Figure 3.**
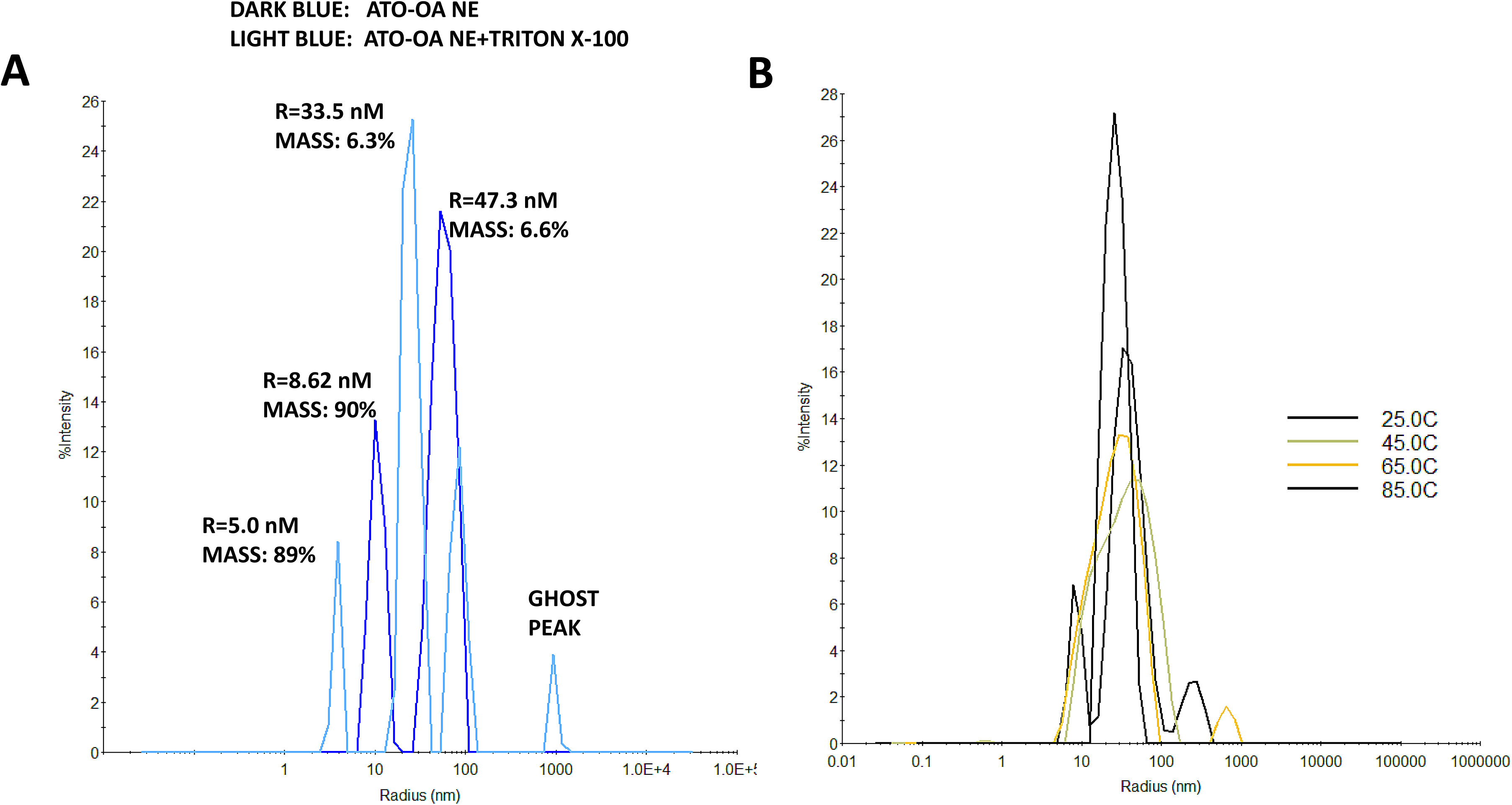
Dynamic light scattering (DLS) analysis of ATO-OA NE. A. Hydrodynamic radius without and with 0.1% Triton-X100. B. Effect of increasing temperature on hydrodynamic radius of ATO-OA NE.

Fig 3B is data showing that ATO-OA NE are thermally stable. There was no significant change in the hydrodynamic radius of ATO-OA NE with heating up to 85°C, suggesting an exceptional thermal stability of ATO-OA NE.

### ATO-OA NE inhibit LAM binding *in vitro*

To assess the efficacy of ATO-OA NE in inhibition of LARP6 binding to 5’SL RNA we added different concentrations of ATO-OA NE to the *in vitro* LARP6/5’SL RNA binding reactions. The concentrations of ATO-OA NE are given as µg/ml. Fig 4 shows that at 3.1 µg/ml ATO-OA NE suppressed LAM binding to A1 5’SL and that at 6.25 µg/ml the binding was almost completely abolished. For comparison, the cefixime preparation from which the ATO-OA NE were purified was completely inactive at low concentrations (Fig 4B).

**Figure 4.**
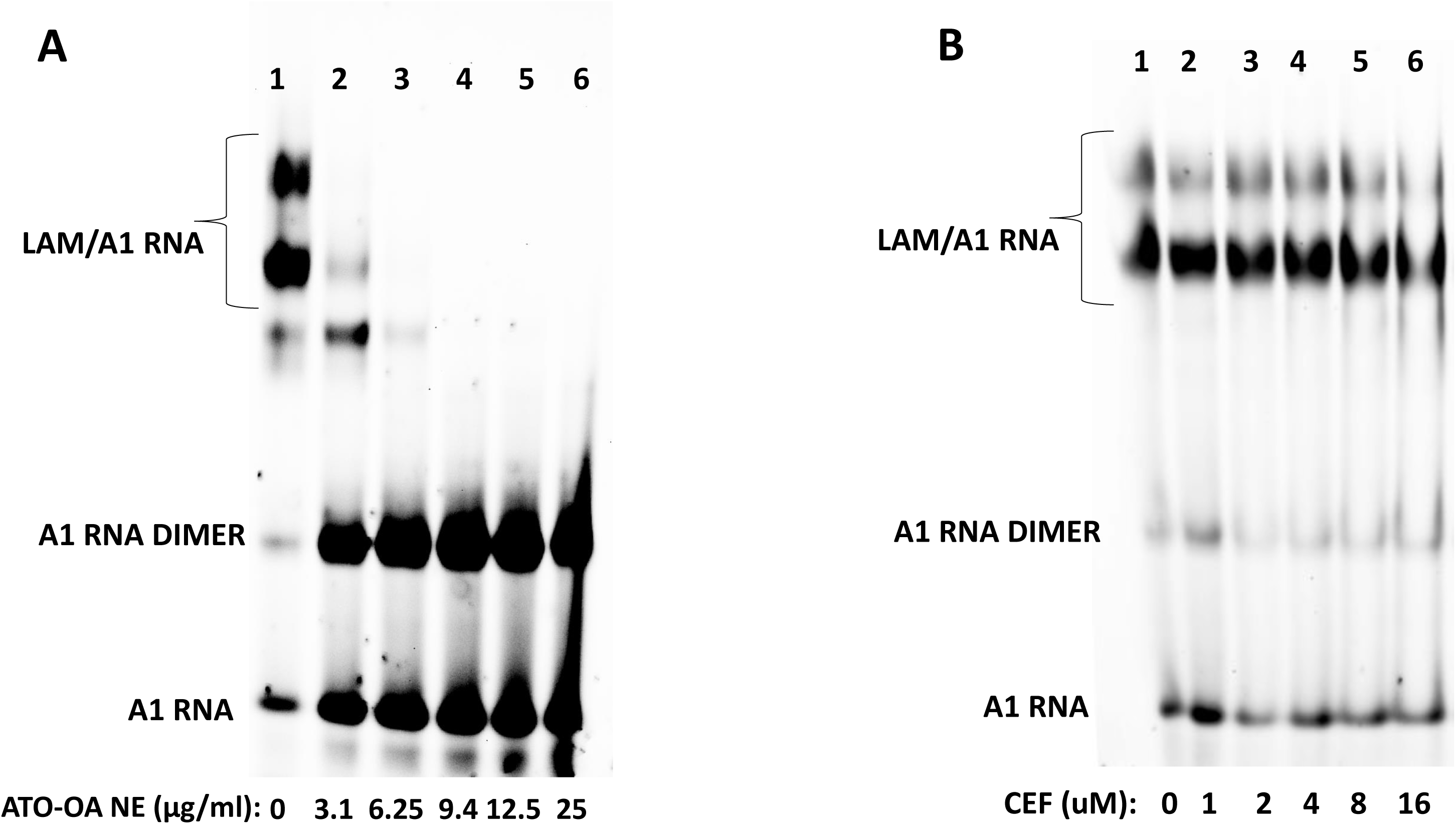
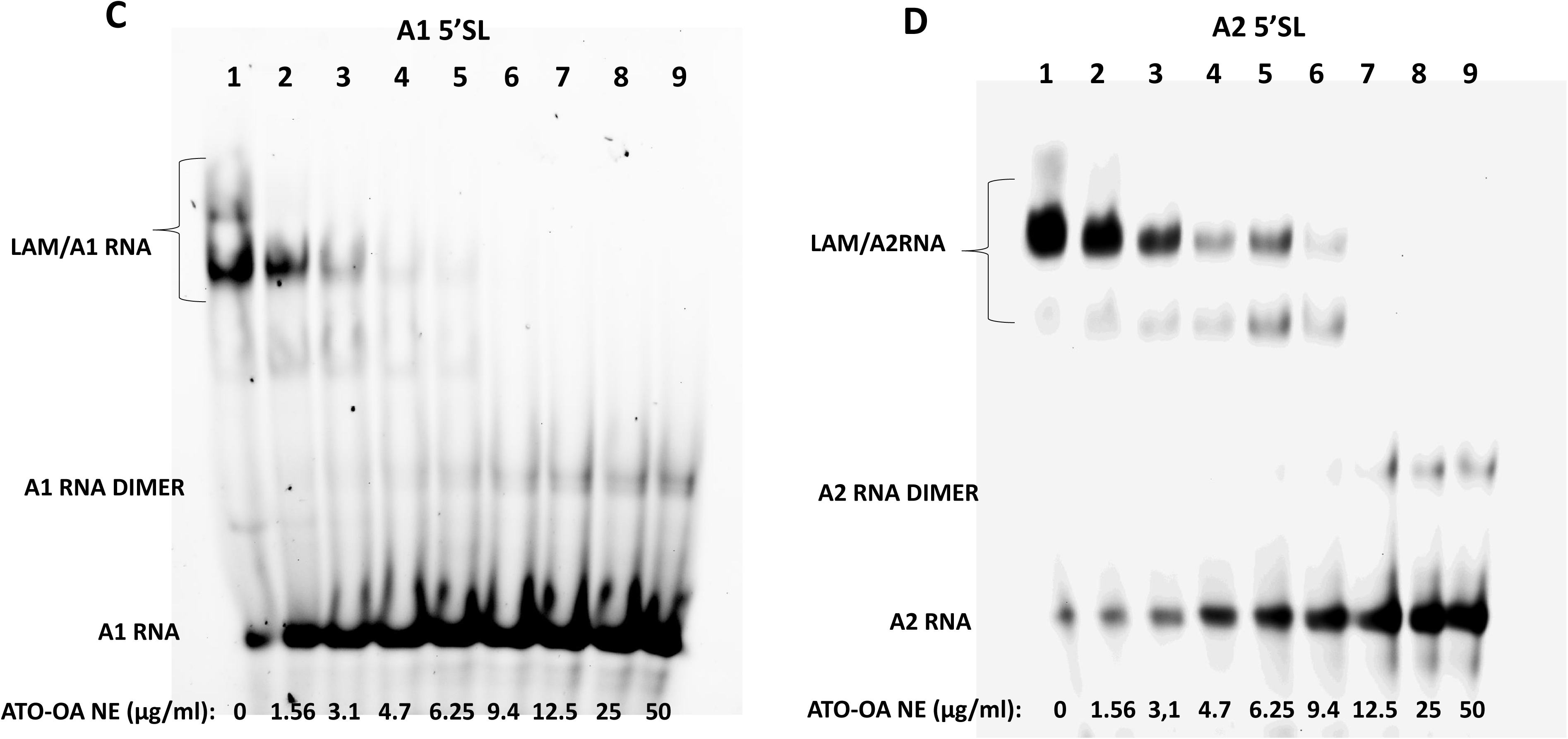
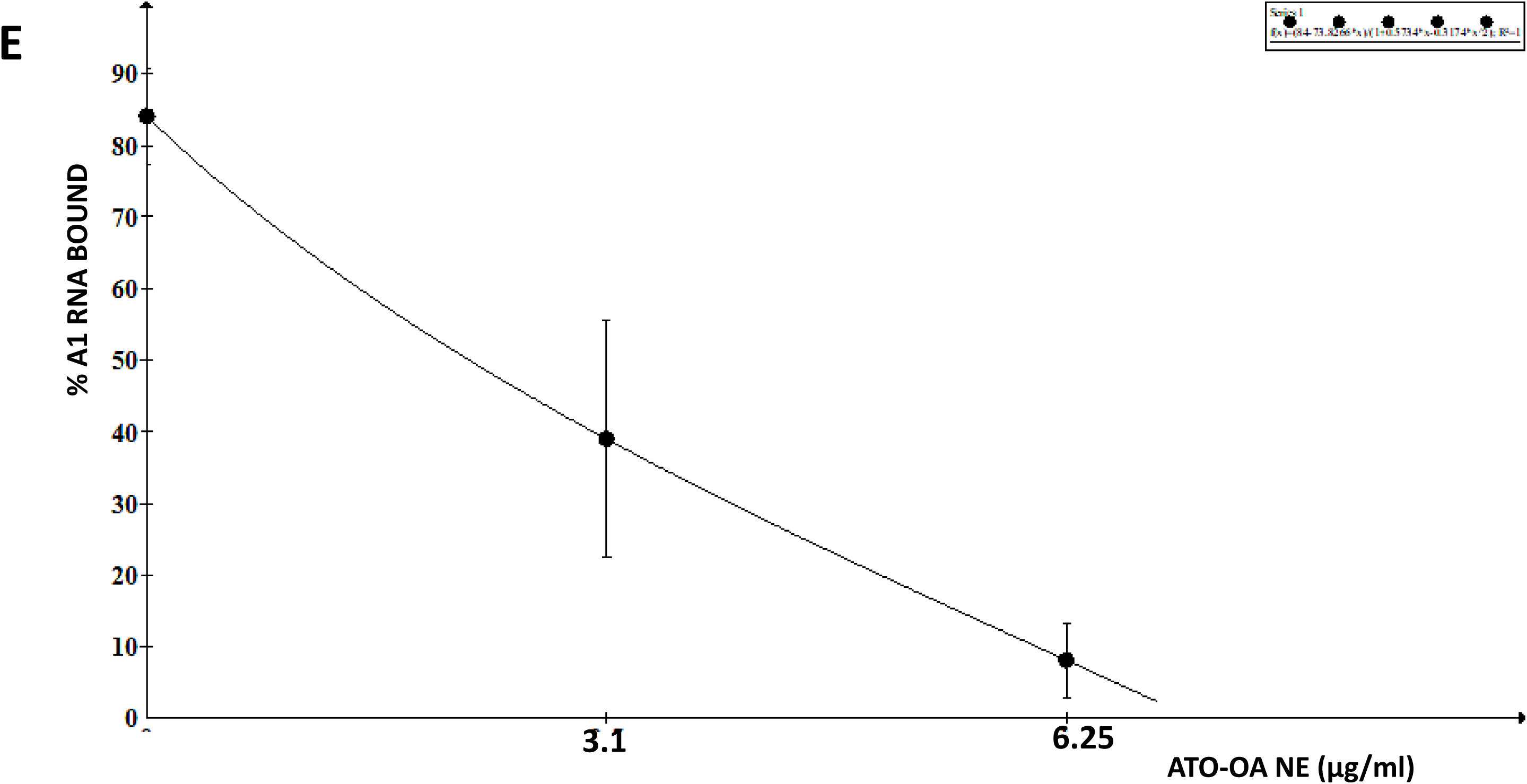
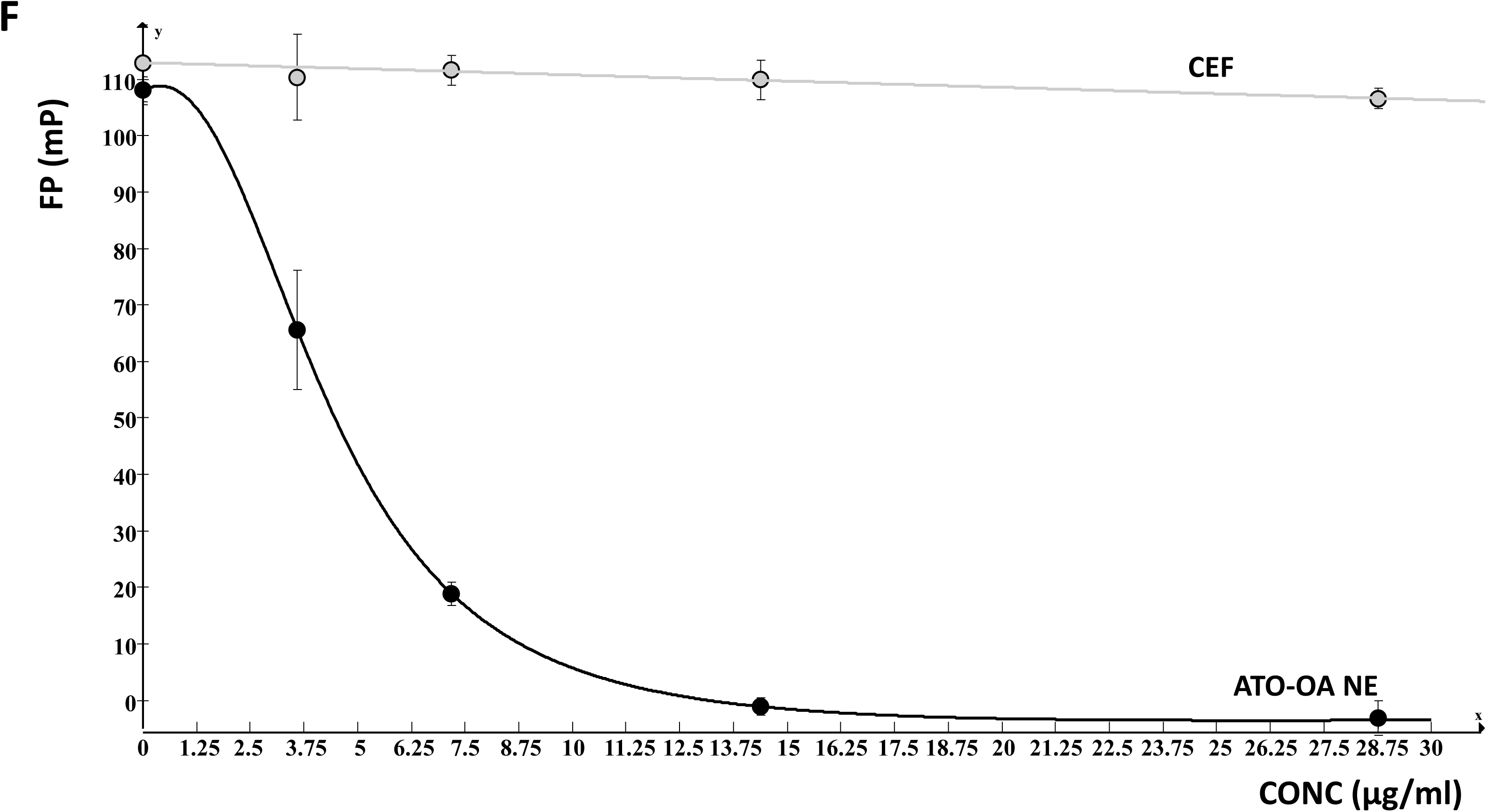
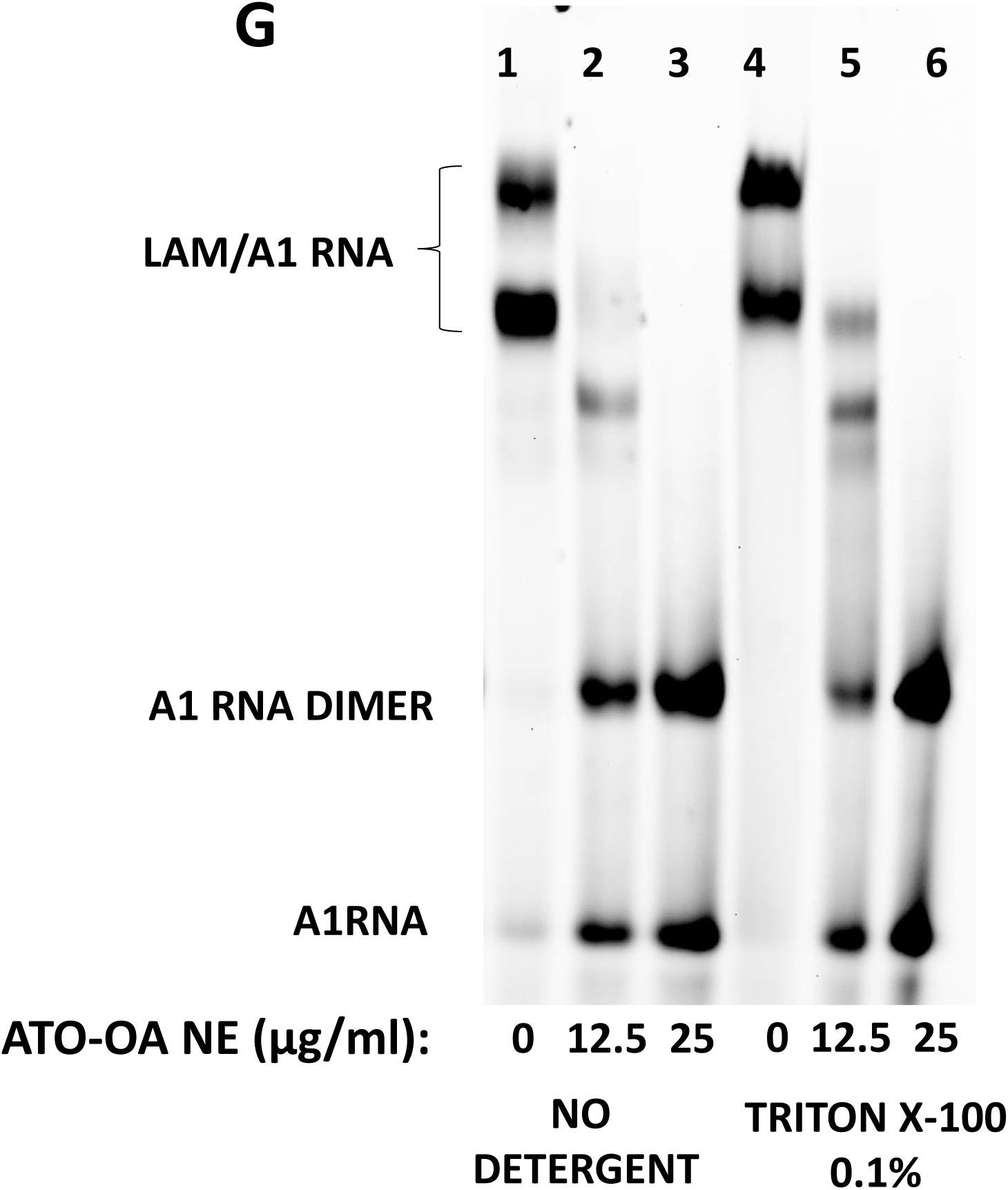
ATO-OA NE inhibits LAM binding. A. Inhibition of LAM binding by ATO-OA NE. Increasing amounts of ATO-OA NE were added to binding reactions and LAM/5’SL RNA complexes and free 5’SL RNA were resolved by gel mobility shift. B. Inhibition of LAM binding by cefixime. C. Titration of ATO-OA NE into A1 5’SL RNA binding reactions. Gel mobility shift experiments with a range of concentrations of ATO-OA NE. Mobility of free RNA monomer (A1 RNA), free RNA dimer (A1 RNA DIMER) and LAM/5’ SL RNA complex (LAM/A1 RNA) is indicated. D. Titration of ATO-OA NE into A2 5’SL RNA binding reactions. Experiment as in C, except A2 5’SL RNA was used. E. Estimation of IC_50_ from gel mobility shift experiments. Signal of A1 RNA in complex with LAM was normalized to the signal of total A1 5SL RNA (% A1 RNA bound) and plotted against ATO-OA NE concentration. Error bars: +-1SD, n=4. F. Dose response inhibition of LAM binding measured by fluorescence polarization (FP). Decrease in FP was plotted as function of ATO-OA NE concentration (µg/ml, black line) or cefixime concentration (gray line). Error bars: +-1SD, n=4. G. Effect of Triton X-100 on ATO-OA NE activity. Lanes 1-3, binding inhibition in absence of Triton X-100, lanes 4-6, binding inhibition in presence of Triton X-100.

To demonstrate that ATO-OA NE could inhibit LARP6 binding to A1 5’SL and A2 5’SL with equal efficacy, we added equal concentrations to the *in vitro* LAM/A1 5’SL and LAM/A2 5’SL binding reactions. Fig 4C and D shows that the inhibition was similar for A1 5’SL binding (4C) and for A2 5’SL binding (4D). About 50% dissociation of LAM/5’SL complexes was seen at 1.56 µg/ml for LAM/A1 5’SL and at 3.1 µg/ml for LAM/A2 5’SL. The gel mobility shift analysis was repeated in four independent replicates, and the intensity of LAM/A1 RNA bands was plotted as function of ATO-OA NE concentration (Fig 4E). From the concentration dependent decay of the complex we estimated the IC50 of ∼3 μg/ml. During the course of these experiments we also observed that, regardless if the LAM/5’SL RNA complex was formed before ATO-OA NE addition or ATO-OA NE were added before complex formation, the inhibition was equally effective.

Fluorescence polarization (FP) is another method to assess LAM/5’ SL RNA binding [48]. In FP experiments we assembled LAM/A1 RNA complex and measured its FP in absence and after addition of increasing concentrations of ATO-OA EN. Decrease in FP indicates the complex dissociation, and the plot of FP vs ATO-OA EN concentration can give an independent estimate of IC_50_. Fig 4F shows that the FP of LAM/A1 RNA complex rapidly diminished with the apparent IC_50_ of ∼4 µg/ml, what was in excellent agreement with the result from gel mobility shift experiments.

The dynamic light scattering measurements with Triton X-100 (Fig 3) suggested that ATO-OA NE are not colloids. To verify that the addition of Triton X100 will not alter the inhibitory potential of ATO-OA NE we added 0.1% Triton X-100 to the *in vitro* binding reactions. Fig 4G shows that ATO-OA NE were equally active in the presence of the detergent.

### ATO-OA NE have no antimicrobial activity

To verify that no antimicrobial activity is associated with the ATO-OA NE we compared the growth of E. coli in presence of cefixime or ATO-OA NE (Fig 5). Cefixime inhibited E. coli growth at concentrations of 0.44 µM and higher, however, ATO-OA NE showed no growth inhibition at lower concentrations, while a small effect at 88 µg/ml could be attributed to a residual cefixime contamination.

**Figure 5.**
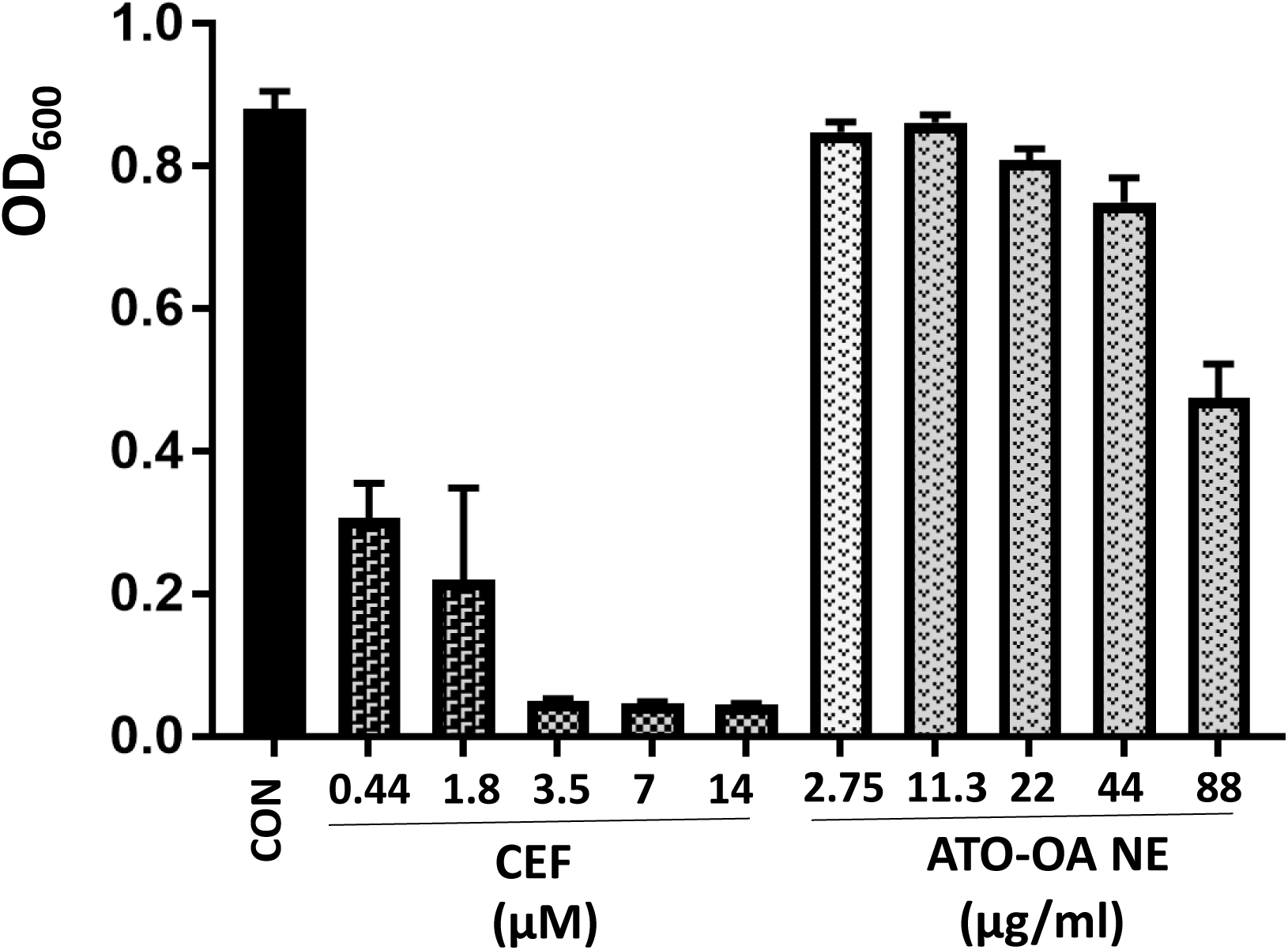
Antimicrobial activity of cefixime and ATO-OA NE. Comparison of E. coli growth inhibition by increasing concentrations of cefixime and ATO-OA NE. CON, no antibiotic.

### Binding of LA domain to 5’SL RNA is not inhibited by ATO-OA NE

LA domain alone can bind 5’SL RNA, albeit with lower affinity then LAM [24, 35]. Therefore, we could test if ATO-OA can inhibit binding of LA domain alone. Fig 6A shows the binding of LA to A1 RNA in presence of increasing concentrations of ATO-OA NE. The complex of LA/5’SL RNA is smaller than that of LAM/5’SL RNA, but it was clearly resolved. There was no attenuation of LA/A1 5’SL complex up to 312 µg/ml of ATO-OA NE (lanes 2-8), with ∼50% decrease at 312 and 625µg/ml (lanes 9 and 10). As a control, LAM/5’SL RNA complex completely disappeared at 12.5 µg/ml of ATO-OA NE (Fig 6B, lane 2). This clearly demonstrated that the LA domain alone cannot be effectively targeted by ATO-OA NE.

**Figure 6.**
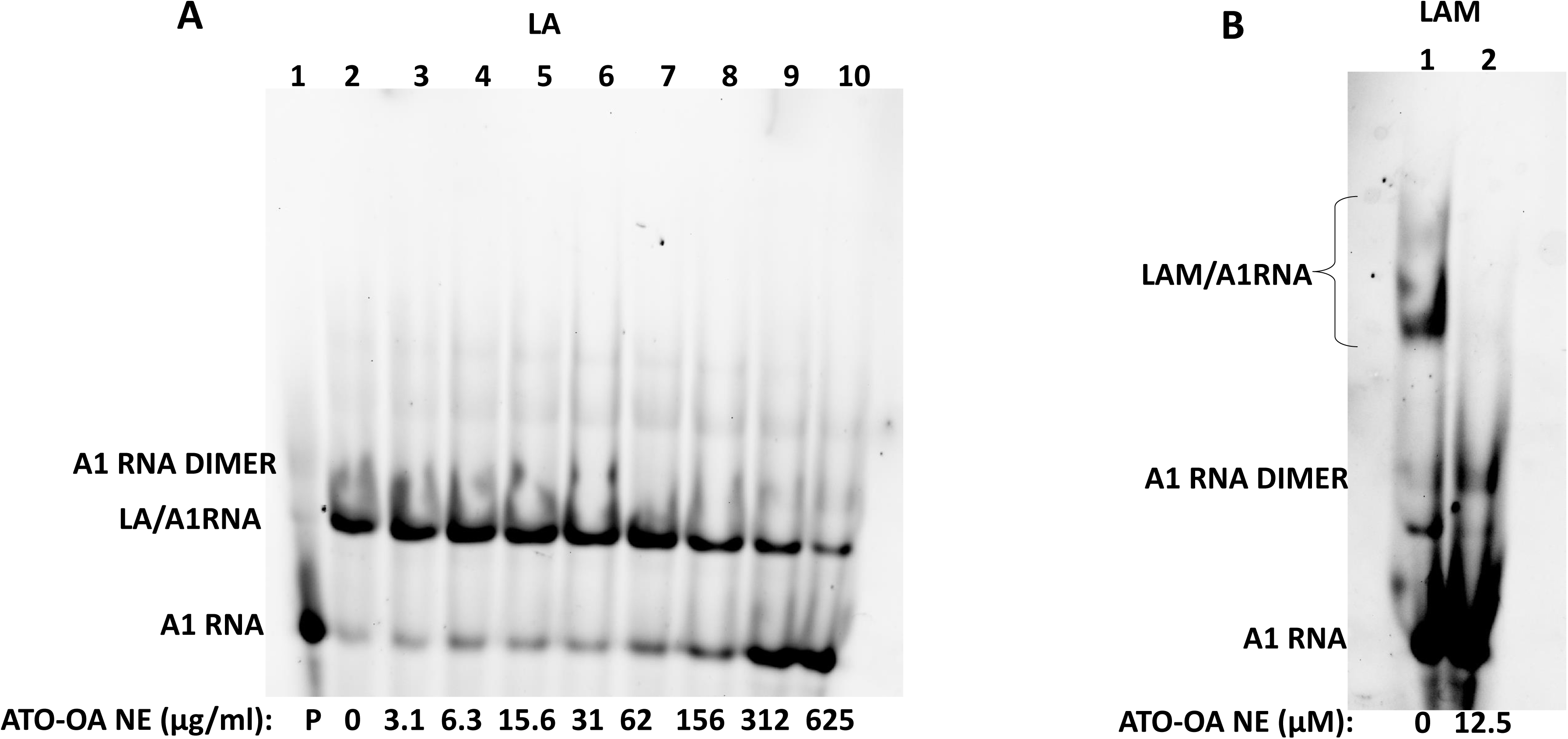
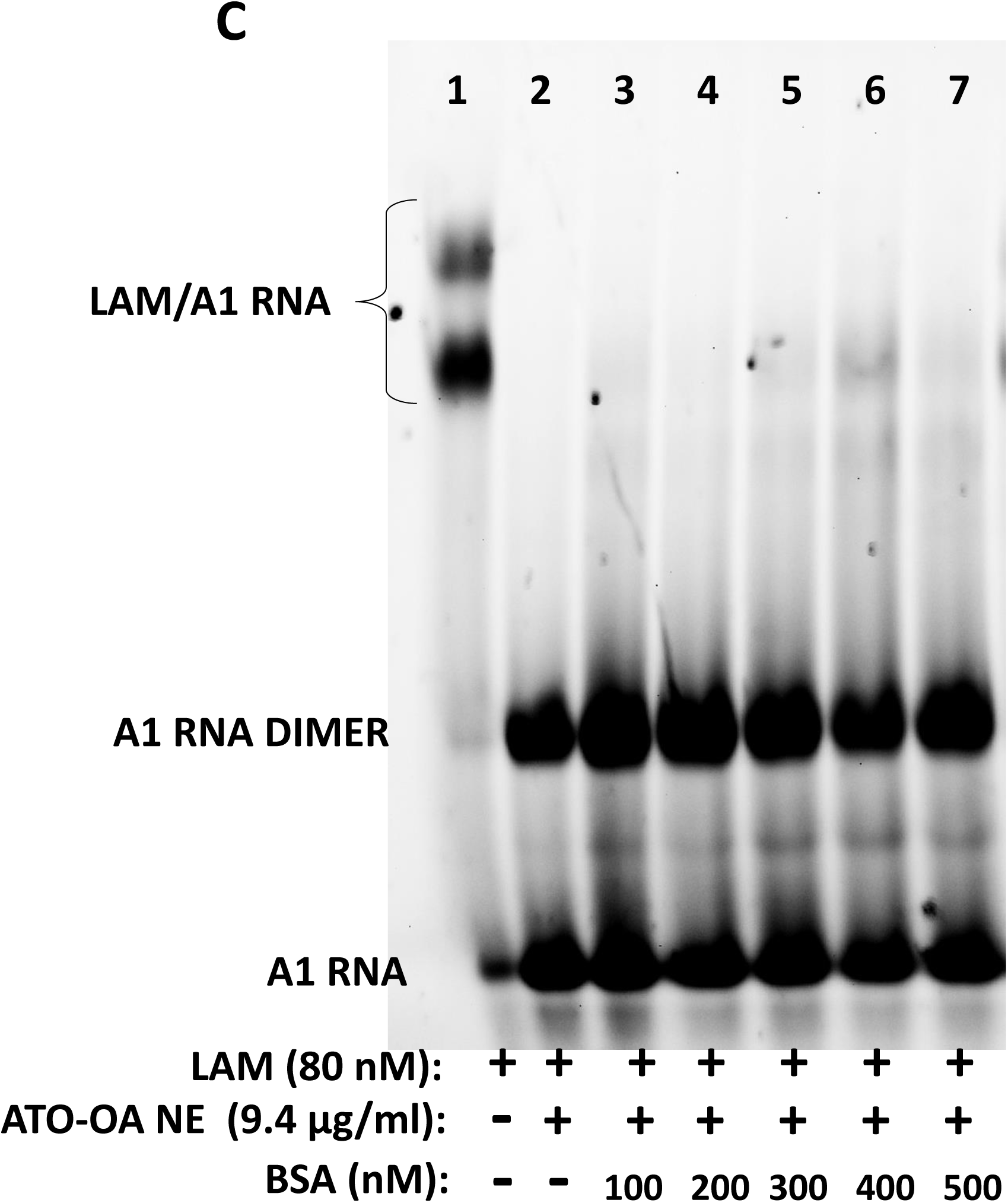
ATO-OA NE has no effect on LA binding. A. Binding of LA to A1 RNA in the presence of increasing concentrations of ATO-OA NE. Mobility of free RNA monomer (A1 RNA), free RNA dimer (A1 RNA DIMER) and LA/5’ SL RNA complex (LA/A1 RNA) is indicated. B. Binding inhibition of LAM. C. BSA does not interfere with ATO-OA NE activity. Inhibition of LAM binding in presence of excess of BSA.

To corroborate that ATO-OA NE is not promiscuous in protein binding we assessed if bovine serum albumin (BSA) could sequester ATO-OA and quench its inhibitory activity for LAM. Fig 6C shows that the presence of 5-fold molar excess of BSA over LAM had no effect on the ATO-OA NE ability to inhibit LAM binding.

### ATO-OA NE interacts with RRM domain of LARP6

To dissociate LAM/5’SL RNA complex ATO-OA NE must interact with LAM or 5’SL RNA or with both. Preliminary experiments showed that ATO-OA NE had no effect on 5’SL RNA, therefore, we prepared recombinant LAM, RRM and LA (Fig 7A) and measured intrinsic tryptophan fluorescence [49] of these proteins (Fig 7B). ATO-OA NE caused quenching of the tryptophane fluorescence of LAM (left panels) and RRM (middle panels), but not that of LA (right panels), as compared to cefixime, which was used as control for inner filter effects. This suggested that ATO-OA NE could alter the conformation of RRM, either alone or as part of LAM.

**Figure 7.**
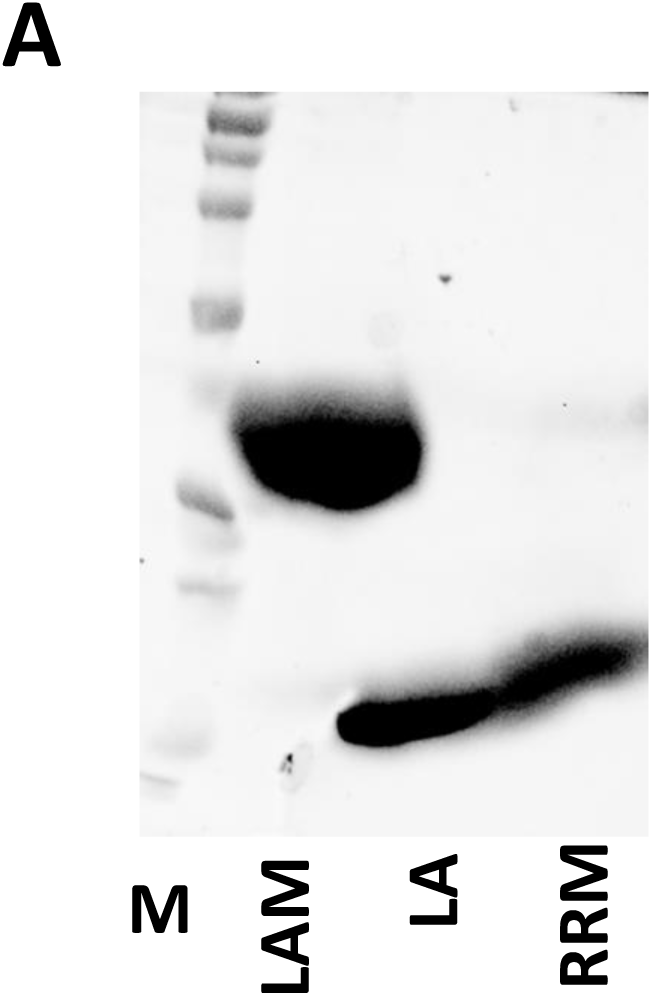
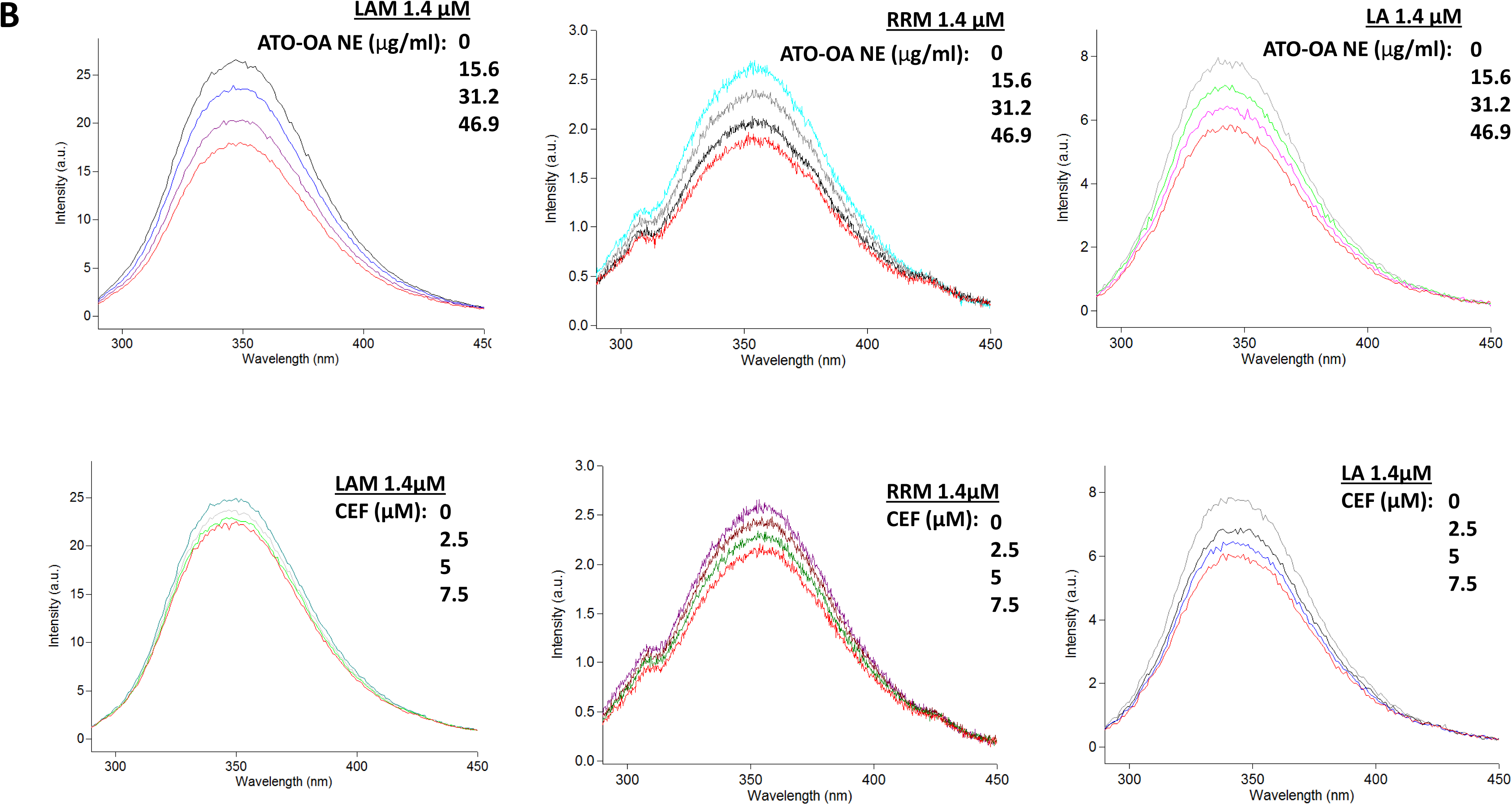
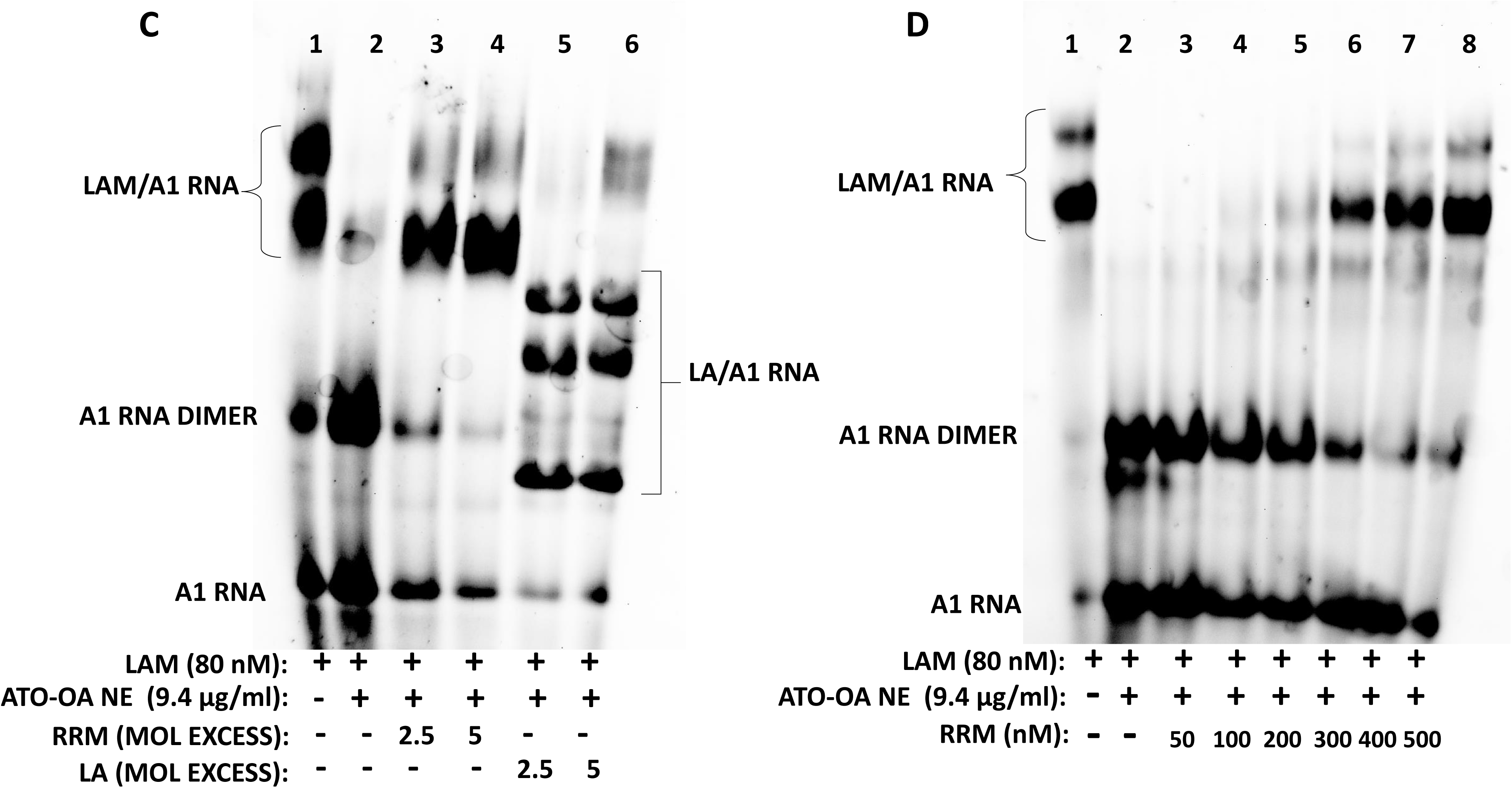
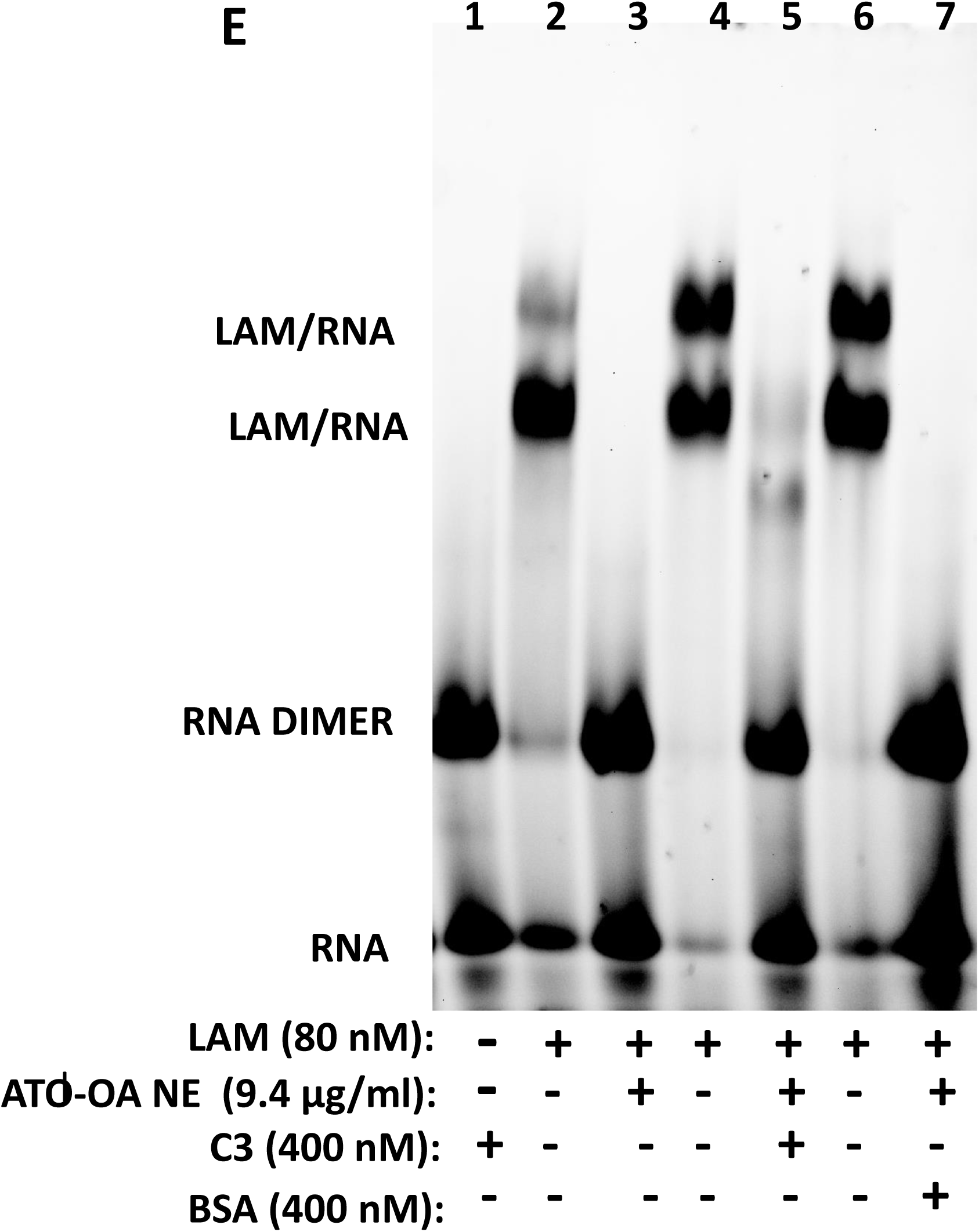
ATO-OA NE induces conformational change of LAM and RRM. A. Proteins used in assays. Coomassie staining of purified LAM, LA and RRM. M, size marker. B. Changes of intrinsic fluorescence of tryptophan. Equimolar amounts of LAM (left panels), RRM (middle panels) and LA (right panels) were titrated with increasing amounts of ATO-OA NE (top panels) or cefixime (bottom panels) and intrinsic tryptophan fluorescence spectrum was plotted as function of concentration. C. RRM competes for ATO-OA NE. Lane 1, LAM/A1 RNA complexes without ATO-OA NE, lane 2, LAM/A1 RNA complexes inhibited by ATO-OA NE, lanes 3 and 4, excess RRM added to the inhibited LAM/A1 RNA complexes, lanes 5 and 6, excess LA added to the inhibited LAM/A1 RNA complexes. LAM/A1 RNA complexes are indicated to the left and LA/A1 RNA complexes to the right. D. Competition with RRM over range of concentrations. Lane 1, LAM/A1 RNA complexes without ATO-OA NE, lane 2, complexes inhibited by ATO-OA NE, lanes 3-8, addition of increasing amounts RRM to the inhibited LAM/A1 RNA complexes. E. Unrelated proteins do not compete for CC28 NE. Lane 1, C3 protease does not bind 5’SL RNA, lanes 2, 4 and 6, binding of LAM, lane 3, inhibition of LAM binding by ATO-OA NE, lane 5, inhibited LAM/RNA complexes in presence of excess C3 protease, lane 7, inhibited LAM/RNA complexes in presence of excess BSA.

### LAM and RRM compete for ATO-OA NE

To further corroborate the interaction of ATO-OA NE with RRM we performed competition experiments between LAM and RRM. LAM was first bound to A1 5’SL RNA (Fig 7C, lane 1), then ATO-OA NE was added to inhibit the binding of LAM (lane 2). When 5 and 10-fold molar excess of RRM was supplemented, the inhibition of LAM binding was relieved and the LAM/A1RNA complex was restored (lanes 3 and 4). This suggested that the excess RRM sequestered ATO-OA NE. The excess of LA domain, which cannot bind ATO-OA NE, did not restore the LAM/5’SL RNA complex, instead LA domain formed its own complexes with 5’SL RNA (lanes 5 and 6). This experiment indicated; first, that the ATO-OA NE inhibition of LAM binding is reversible and, second, that RRM can sequester the inhibitor.

Fig 7D shows titration with RRM over wide range of concentrations. The first hint of RRM rescuing LAM binding was seen at their equimolar concentrations (lane 4), when a tracing amount of LAM/A1 RNA complex reappeared. Almost complete recovery of LA/A1 RNA complex was seen with 5-fold excess of RRM.

To verify that unrelated proteins cannot compete for ATO-OA NE, Fig 7E shows that C3 protease (lane 5) or BSA (lane 7) could not restore LAM binding. This strongly suggested that RRM can sequester ATO-OA NE while unrelated proteins cannot.

### Inhibition of type I procollagen secretion by hepatic stellate cells (HSCs) in culture

HSCs are cells responsible for liver fibrosis [50, 51] and here we used LX-2 human HSCs line [52] to test the effects of ATO-OA NE. Fig 8A shows that when HSCs were incubated for 24h with ATO-OA NE, their ability to secrete type I procollagen (COL1A1) was reduced at concentrations of 125 µg/ml and 250 µg/ml and degradation fragments of COL1A1 polypeptide appeared in the cell medium (COL1A1 DEG). This indicated that *in vivo* ATO-OA NE impaired productive assembly and secretion of type I collagen.

**Figure 8.**
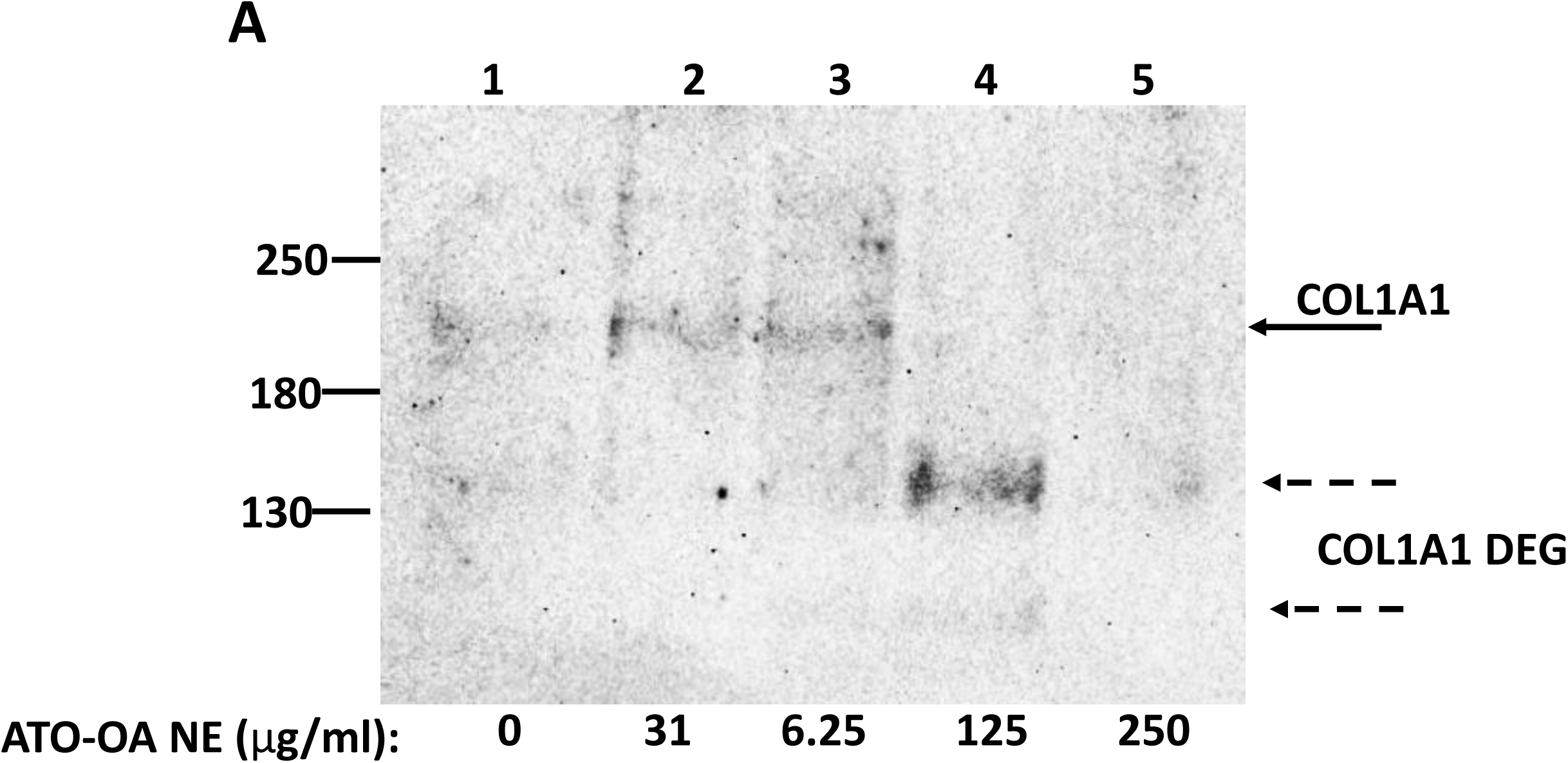
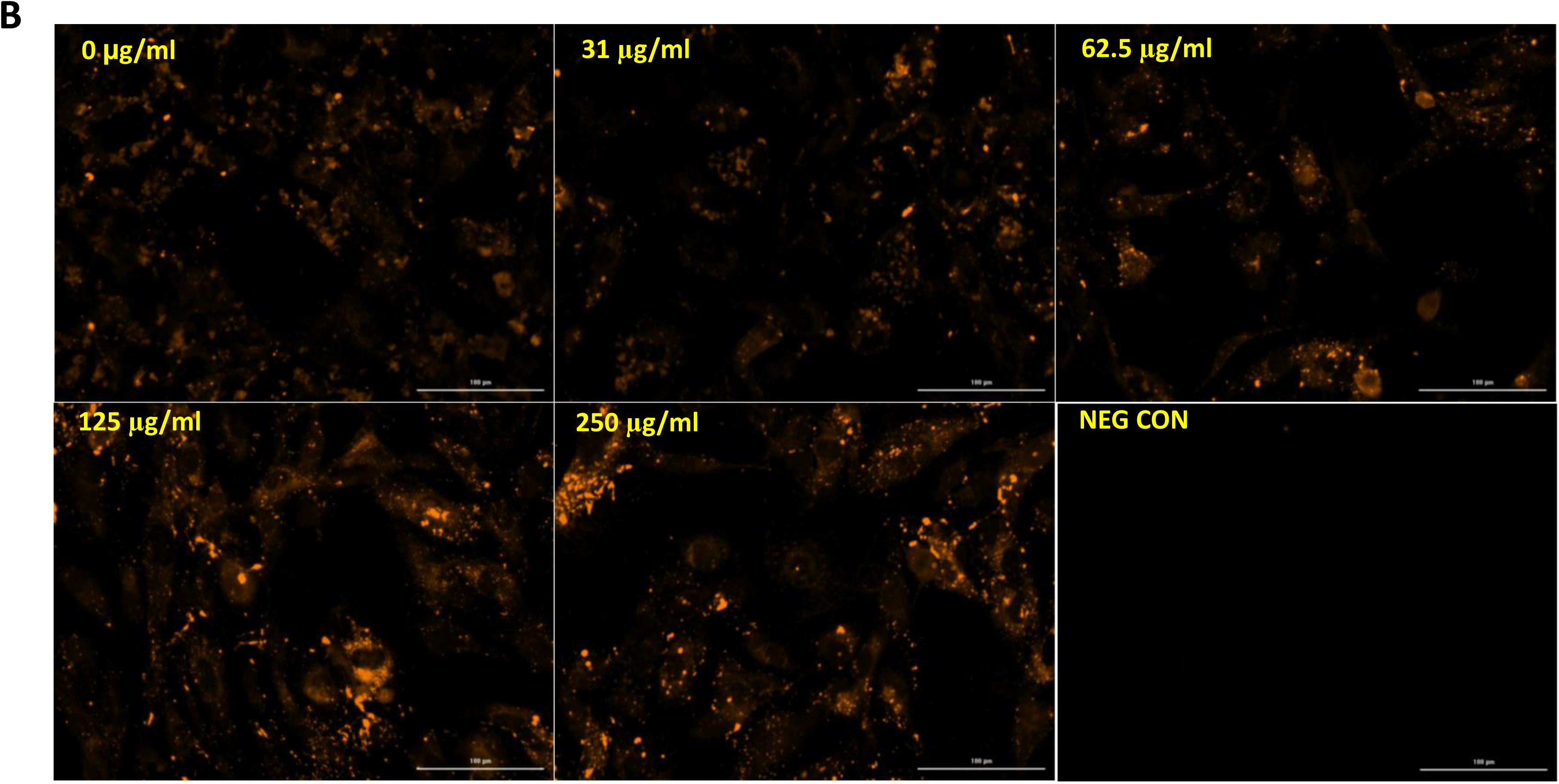
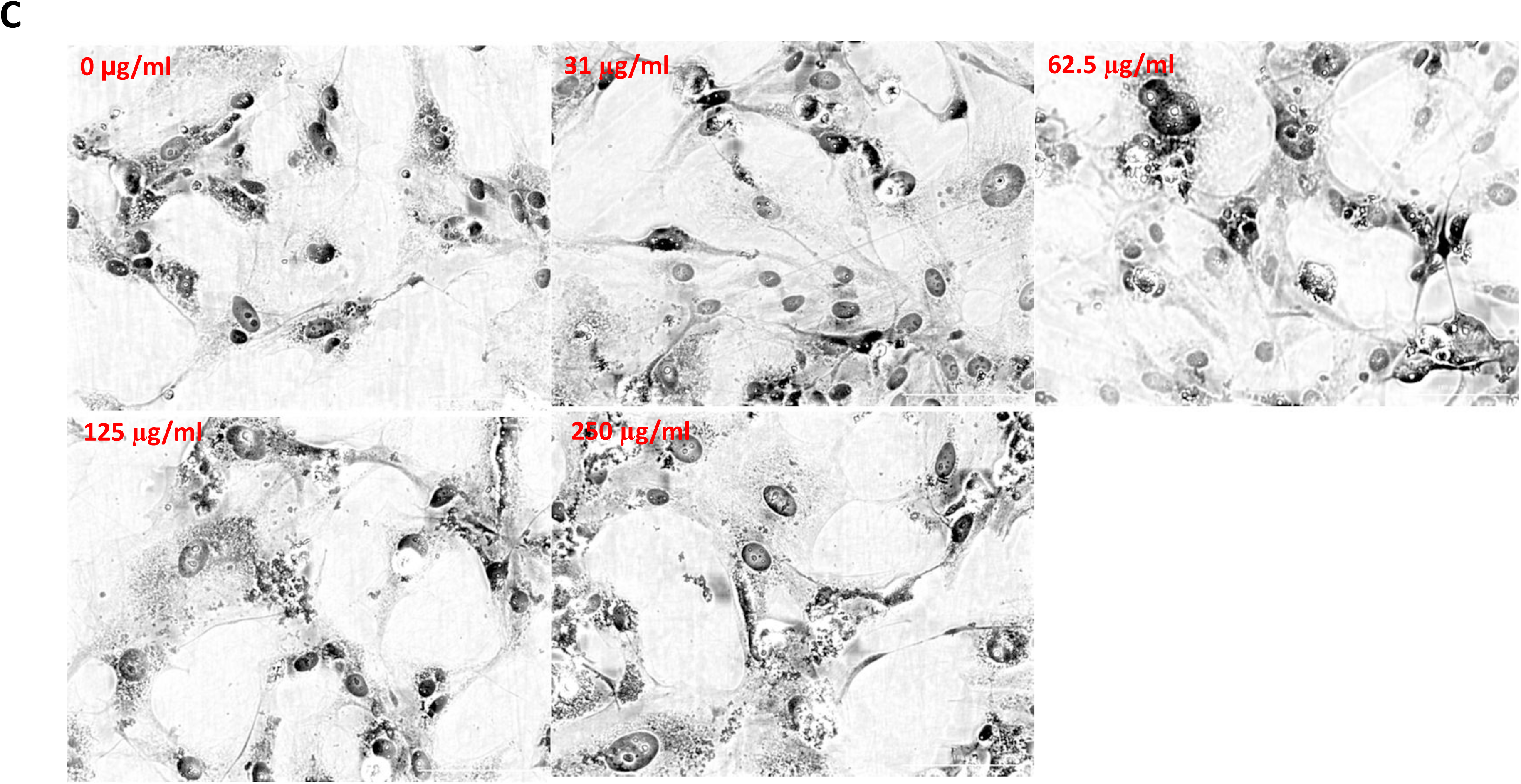
Effect of ATO-OA NE on type I procollagen secretion by HSCs. A. Human HSCs were treated with the indicated concentrations of ATO-OA NE for 48h and COL1A1 polypeptide was measured in cellular medium by western blot. Full size COL1A1 peptide and its degradation products are indicated. B. Intracellular accumulation of type I procollagen. HSCs were treated with indicated concentrations of ATO-OA NE and intracellular type I procollagen was visualized by immunostaining (20x magnification). NEG CON, immunostaining without anti-collagen antibody. C. Phase contrast images of HSCs treated with ATO-OA NE.

Next, we assessed if the poor secretion was associated with excessive intracellular retention of procollagen. The intracellular procollagen was assessed by immunostaining of HSCs using type I collagen specific antibody (Fig 8B). In control cells the intracellular type I procollagen showed faint immunostaining in the perinuclear region of the cells (0 µg/ml). This pattern remained unchanged with 31 µg/ml and 62.5 µg/ml of ATO-OA NE. However, when the cells were treated with 125 µg/ml and 250 µg/ml of ATO-OA NE, intense intracellular staining and presence of larger aggregates of type I procollagen was seen. Quantification of the number of high intensity pixels showed about 2-fold increase with 125 µg/ml of CC297 NE (p=0.0915) and 3.3-fold increase with 250 µg/ml of ATO-OA NE (p=0.0189). This result indicated that the ATO-OA NE impaired secretion of type I procollagen was associated with its cellular retention. The morphology of LX-2 cells and their number was not changed by treatment with ATO-OA NE (Fig 8C).

### Inhibition of type I procollagen secretion by human lung fibroblasts (HLFs) in culture

HLFs are cells responsible for pulmonary fibrosis [53]. When HLFs were incubated with ATO-OA NE, the intracellular retention of type I procollagen was seen at 125 µg/ml and 250 µg/ml of ATO-OA NE, the same concentrations that had the effect in HSCs (Fig 9A). Overlaying the collagen staining with the phase contrast image of cells revealed that type I procollagen accumulated in the perinuclear regions as intensely stained granules (Fig 9B).

**Figure 9.**
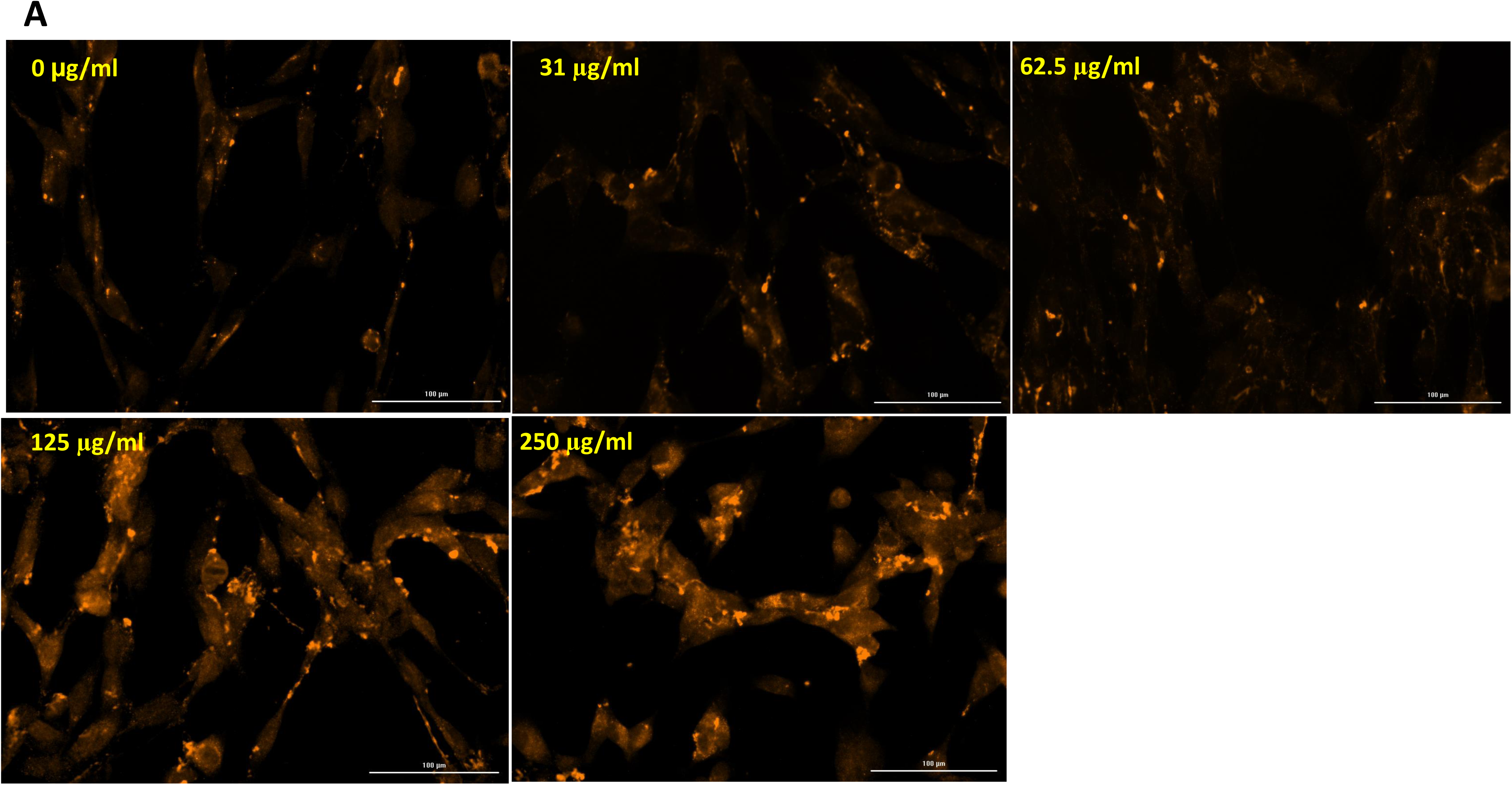
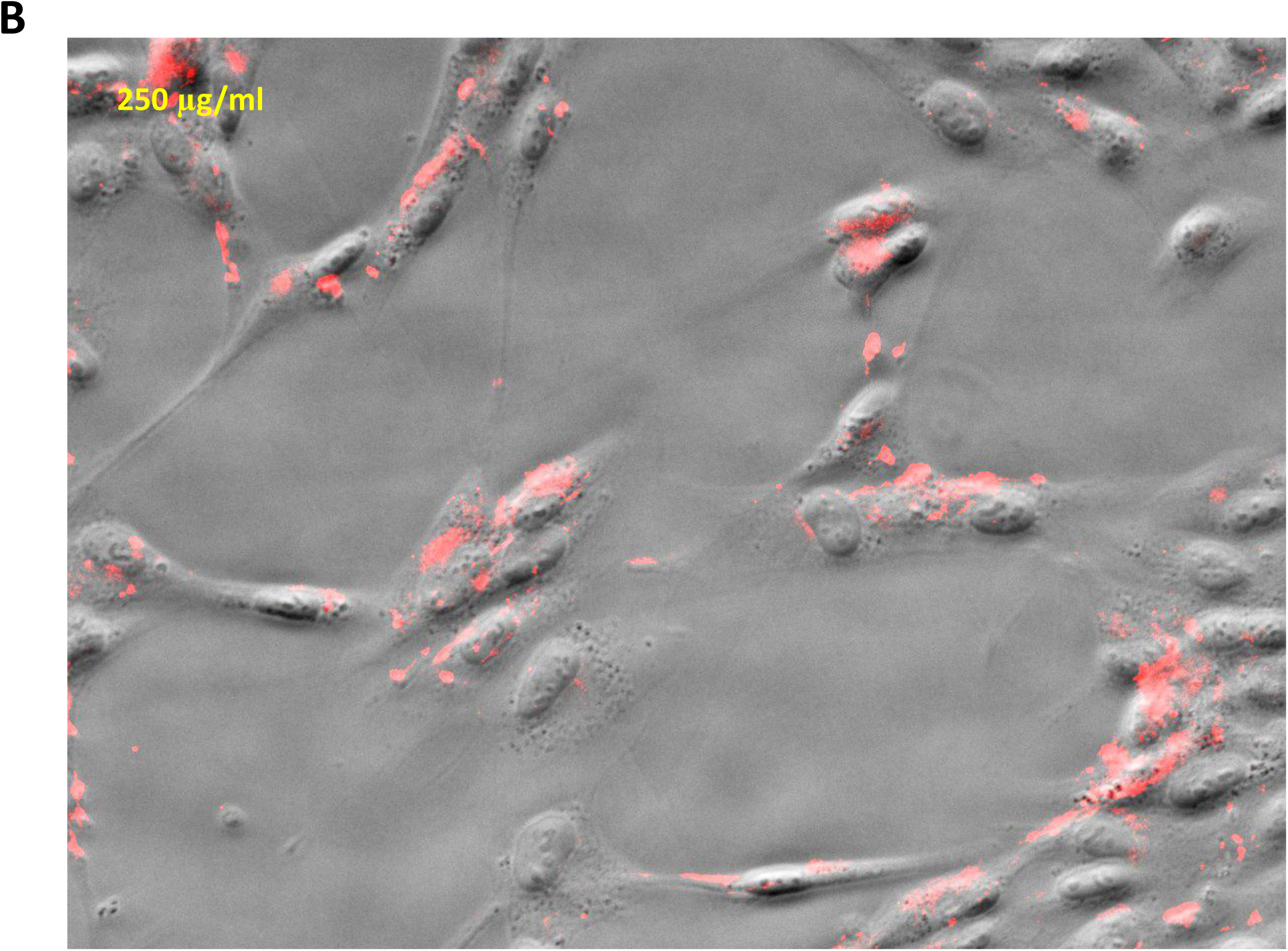
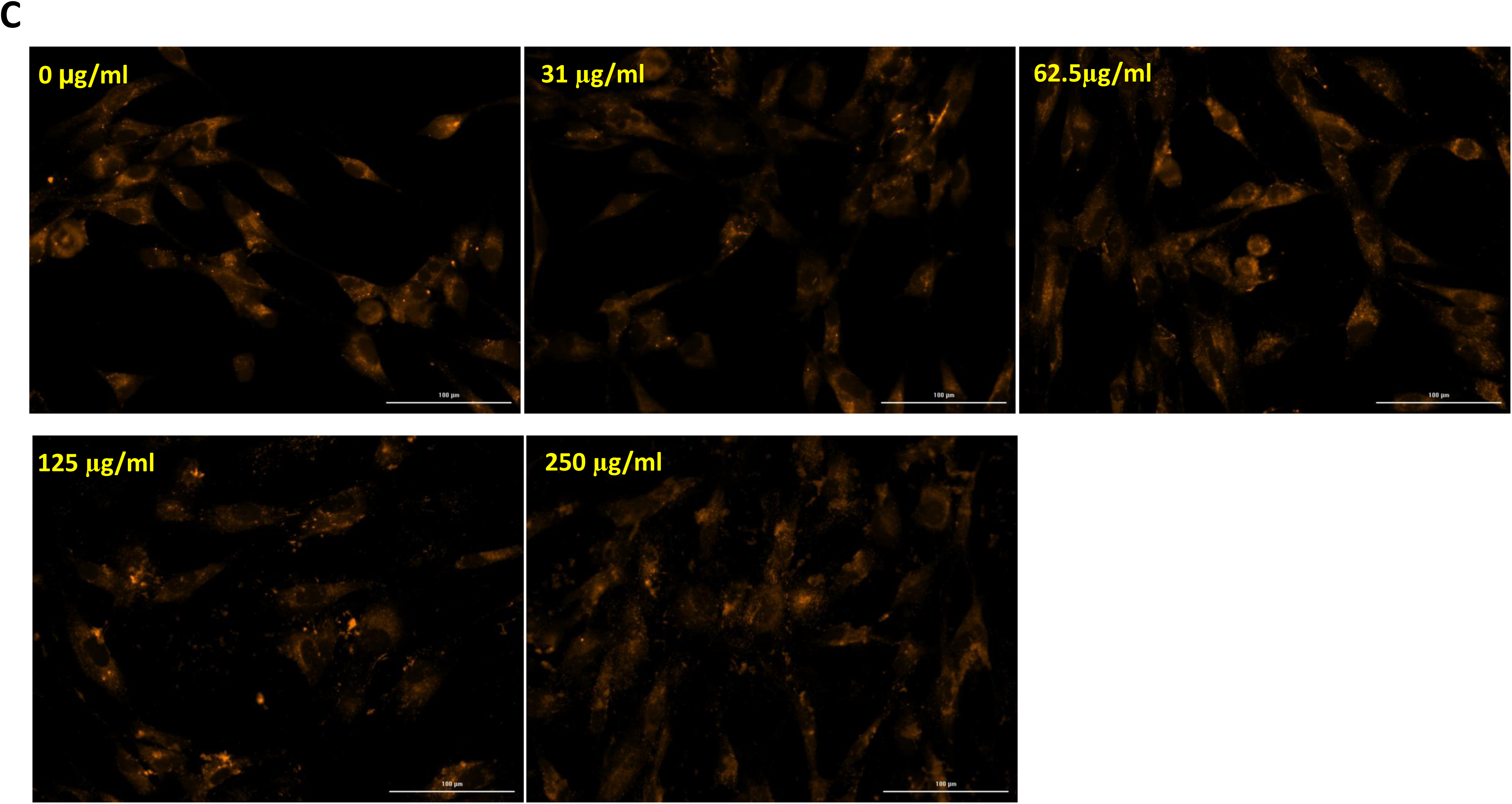
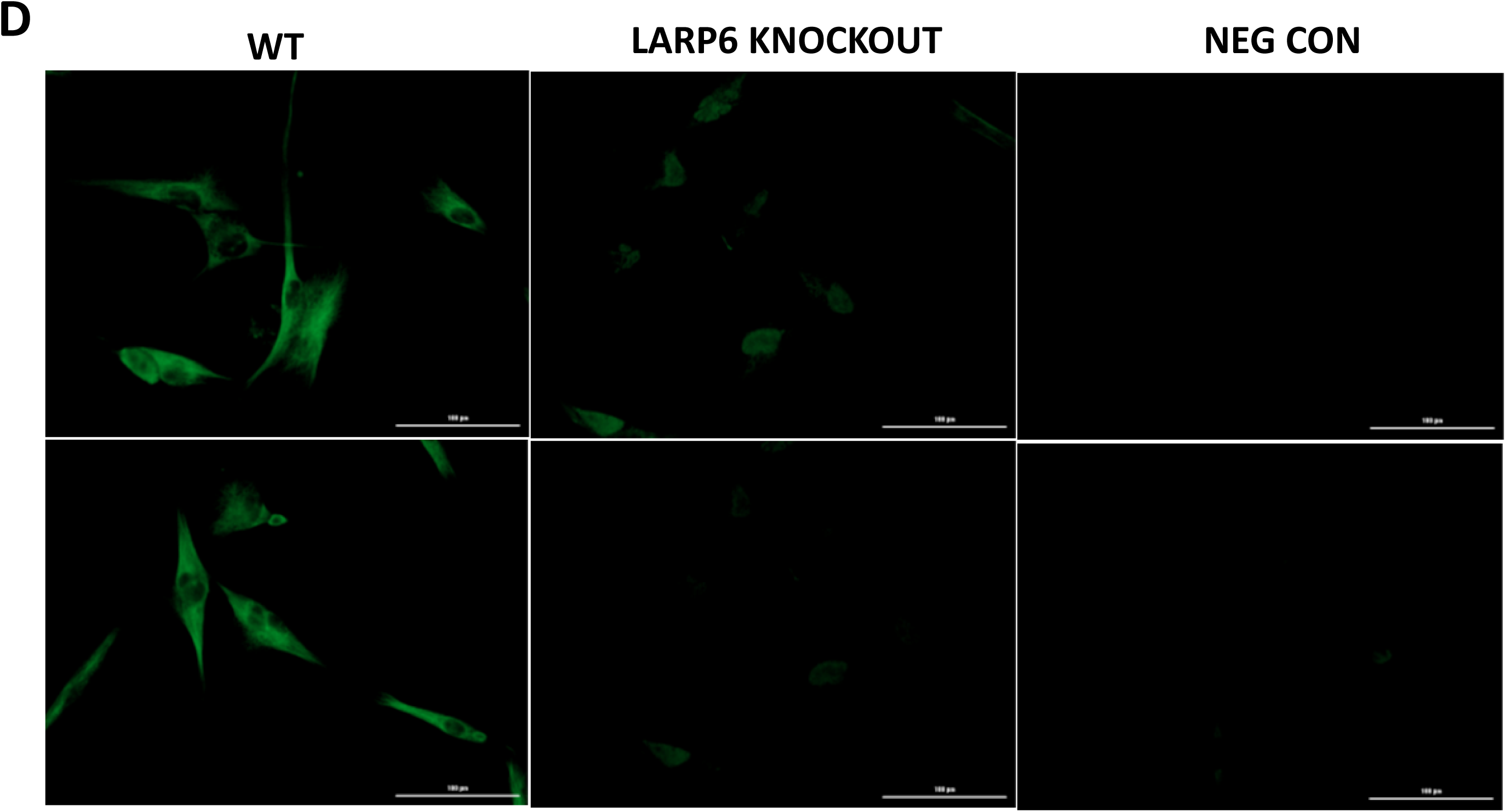
Intracellular accumulation of type I procollagen in HLFs after ATO-OA NE treatment. A. Immunostaining of type I procollagen in HLFs after treatment with indicated concentrations of ATO-OA NE (20x magnification). B. Overlay of procollagen staining and phase contrast image of HLFs. C. Effect of ATO-OA NE on intracellular type I procollagen accumulation in LARP6 knockout HLFs by CRISPR/CAS9. D. Expression of LARP6 in WT and CRISPR/CAS9 knockout HLFs. LARP6 was visualized by immunostaining. NEG CON, immunostaining without anti-LARP6 antibody (20x magnification).

To investigate if the effect of ATO-OA NE depends on LARP6 we knocked out LARP6 in HLFs by CRISPR-CAS9 and incubated the knockout cells with ATO-OA NE. Collagen immunostaining showed no difference in the intracellular accumulation of type I procollagen between the knockout cells treated with or without ATO-OA NE (Fig 9C), suggesting that the inhibitory effect of ATO-OA NE depends on presence of LARP6. Fig 9D shows the expression of LARP6 in WT and knockout HLFs.

### Fibroblasts from 5’SL knock-in mice do not respond to ATO-OA NE

The knock-in mice were created where a mutation of 5’SL sequence in COL1A1 gene was made, but without altering the expression or coding region of the gene [31]. This gene encodes for COL1A1 mRNA without 5’SL and which cannot be regulated by binding LARP6. The knock-in mice are viable but are resistant to development of hepatic fibrosis or fibrosis of arterial cell walls [27, 31]. Mouse embryonic fibroblasts (MEFs) were derived from the WT littermates (WT) and homozygous knock-in mice (5’SL knock-in). These cells provided an additional opportunity to test if the effect of ATO-OA NE on type I procollagen production is dependent on LARP6 binding. Fig 10 shows type I collagen immunostaining in WT MEFs and 5’SL knock-in MEFs treated with 125 µg/ml of ATO-OA NE. While there was increased intracellular accumulation of type I procollagen in WT MEFs after ATO-OA NE treatment, there was no change in the 5’SL knock-in MEFs. This suggested that ATO-OA NE is effective only in cells which synthesize type I collagen in the LARP6 dependent manner.

**Figure 10.**
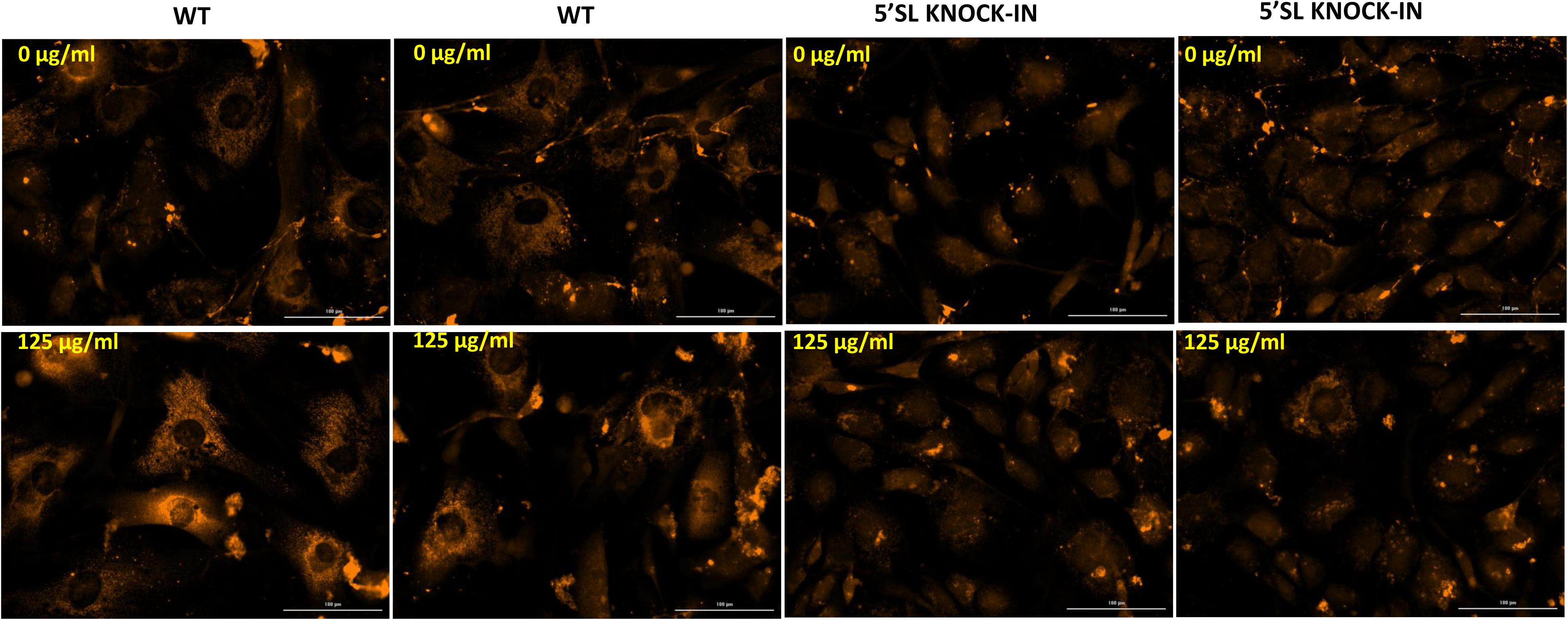
Knock-in MEFs with mutation of 5’SL of COL1A1 mRNA do not respond to ATO-OA NE. Immunostaining of type I procollagen in WT MEFs and 5’SL knock-in MEFs after treatment with 125 µg/ml of ATO-OA NE (20x magnification).

### ATO-OA NE decreases type I procollagen production by organoids in culture

To assess the effect of ATO-OA NE on type I procollagen biosynthesis in organoids, the organoids of human pancreatic adenocarcinoma cells were grown in three dimensions matrix. These cells form organoids of different sizes after three days of growth in matrigel [54]. We treated the organoids with 62.5 µg/ml of ATO-OA NE during their growth and immunostained the organoids at day 3 for type I collagen and actin. Actin was used to visualize the organoids and to normalize collagen expression. Confocal images of multiple organoids were taken at 20x magnification, arranged in stacks and the maximal projection of stacks is shown in Fig 11. Actin staining was in green and collagen staining was in red, so each green stained object represents an organoid with type I collagen superimposed. Control organoids showed foci of strong red staining, revealing robust type I collagen accumulation at discrete sites (Fig 11C, upper panels). In treated organoids the foci were mostly absent, and collagen staining was weak and diffuse (lower panels). By counting high intensity red pixels and normalizing them to the number of green pixels we estimated about 3-fold reduction in type I procollagen accumulation (p=0.0197, n=3).

**Figure 11.**
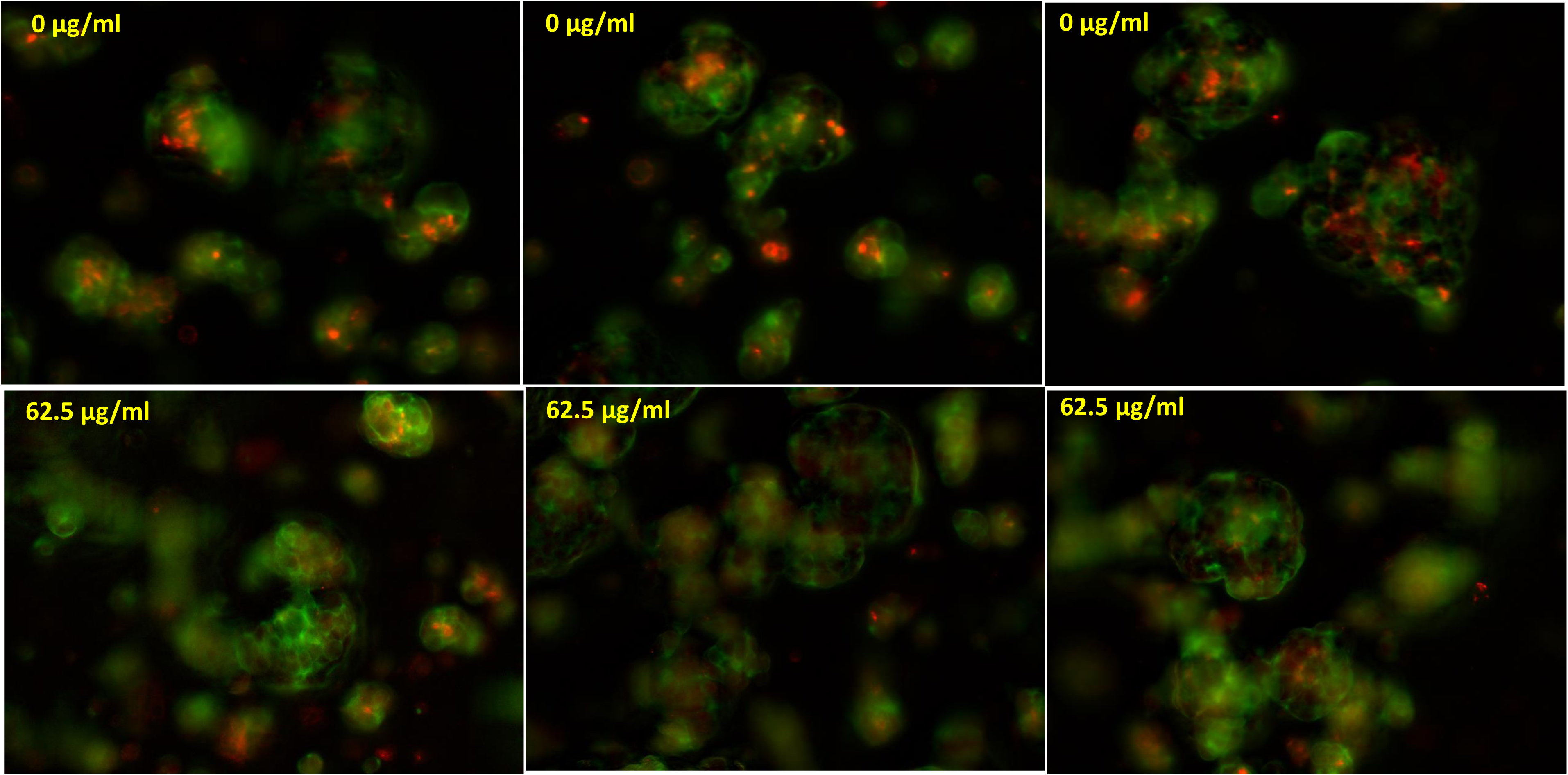
ATO-OA NE suppresses type I procollagen production by human pancreatic adenocarcinoma organoids. Images of organoids (green, actin staining) with superimposed type I procollagen staining (red) in untreated organoids (upper images) and organoids treated with 62.5 µg/ml of ATO-OA NE for 3 days (20x magnification). Maximal projection of confocal stacks of three viewing fields is presented.

## Discussion

Organ fibrosis is a chronic, progressive disease that requires prolonged treatment. Ideal antifibrotic drugs must have minimal side effects, be affordable and suitable for years of therapy. Chemical compounds that can specifically suppress persistent type I collagen biosynthesis are, therefore, desirable. The discovery that binding of LARP6 to 5’SL sequence of type I collagen mRNAs is a key regulatory step of type I collagen production in fibrosis [27, 31] prompt the search for LARP6 binding inhibitors [32, 41]. Here we describe the finding that some commercial preparations of the third-generation cephalosporin, cefixime, contained an inhibitory activity for LARP6 binding to 5’SL. The inhibitor was identified as ion with [M+Z]^+^ of 287. This compound was in the form of nano-entities (NE) when extracted from cefixime. It was termed ATO-OA NE and the results described here were obtained using the ATO-OA NE purified from cefixime.

In this manuscript we show that: 1. Nano-entities (NE) formed by ATO-OA are critical for activity, but they are not colloidal aggregates and have a mean hydrodynamic radius of 1.9 nM. 2. ATO-OA NE inhibit formation of LAM/5’SL RNA complexes *in vitro* with IC_50_ of 3-4 μg/ml. 3. ATO-OA NE interact with the RRM domain of LARP6 and alter its conformation; the induced allosteric change converts the domain from an enhancer of LARP6 binding into a dominant negative domain for LARP6 binding. 4. ATO-OA NE bind the RRM of LARP6 specifically and are not promiscuous protein binders. 5. Treatment of HSCs and HLFs with ATO-OA NE results in poor secretion of type I procollagen and its intracellular retention. 6. This effect is dependent on the presence or activity of LARP6 in the cells. 7. tissue organoids treated with ATO-OA NE have reduced type I collagen production. These findings provide a strong premise that the compounds based on ATO-OA NE scaffold can be developed into effective and specific antifibrotic drugs.

Many FDA approved drugs can self-associate into nano-entities having sizes of 1-100 nM. Most drugs form colloidal nano-entities, which is an undesired property regarding drug screening [55]. Colloidal particles cause false positive hits in drug screening efforts, as they tend to nonspecifically associate with proteins and inhibit their activity *in vitro* [56]. ATO-OA NE are not colloidal aggregates, because their size (Fig 3A) or biological activity (Fig 4G) cannot be abrogated by detergents, such as Triton X-100. ATO-OA NE can withstand heating up to 85°C (Fig 3B), indicating that they are stable. ATO-OA NE selectively associate with the RRM domain of LARP6 (Fig 7) and not with the LA domain, BSA or C3 protein, suggesting that they are not indiscriminate protein binders.

Interaction of ATO-OA NE with the RRM domain of LARP6 induces a conformational change of RRM that can be detected as quenching of the intrinsic tryptophan fluorescence [49]. La-motif of LARP6 (LAM) contains LA domain in tandem with RRM and also shows tryptophan fluorescence quenching with and ATO-OA, while LA domain does not (Fig 7B). So, it is safe to conclude that ATO-OA NE target the RRM domain of LARP6 and that the allosteric alteration caused by ATO-OA NE inactivates LARP6 binding to 5’SL.

How do the changes of RRM conformation inactivate LARP6 binding? The explanation may be in the peculiar mode of recognition of 5’SL RNA by LARP6. LA domain recognizes the noncanonical base pairs which exist within the 5’SL bulge and which create a unique surface where the initial contacts between LA and 5’SL are made [35]. This initial recognition causes an induced fit of LA domain, resulting in more extensive contacts. The induced fit of LA appears to be facilitated and supported by RRM and a stable 5’SL RNA/LARP6 complex is formed [23]. If the conformation of RRM domain is altered in a way that it cannot support the induced fit of LA, the binding can be compromised. Because RRM does not directly contact 5’SL RNA [35], the interaction with ATO-OA NE is possible in free, as well as in bound LARP6. This may be of significance for activity in cells, where LARP6 is found bound to collagen mRNAs, causing dissociation, as well as free nucleo-cytoplasmic shuttling protein [57], causing lack of recognition of collagen mRNAs.

The IC_50_ of ATO-OA NE *in vitro* was estimated as 3-4 µg/ml (Fig 4 E and F). It is difficult to translate this into molar concentration because of large and variable sizes of NE. The average hydrodynamic radius of the ATO-OA NE which still shows full activity is 5.0 nM (ATO-OA NE in presence of 0.1% Triton X-100, Fig 3). This hydrodynamic radius corresponds to the size of a globular protein of ∼99 kD. ATO-OA NE are not proteinaceous, and their shape is unknown; they may be globular but hollow. Therefore, their hydrodynamic radius could not be related to the molecular weight.

It has been assumed that RNA/protein interactions are difficult to disrupt by small molecules [58]. Most small molecules discovered as inhibitors of protein/RNA interactions are active at concentrations between 2.5 µM and 100 µM (LIN-28 inhibitors: 2.5-10 μM [59], HUR inhibitors: 100 μM [60], RBM39 inhibitors: 6-7 μM [61], NONO inhibitor: 5.7 μM [62]). The exceptional potency of ATO-OA NE can be attributable to the size of particles, which may have large surface area to alter the RRM of LARP6. If ATO-OA NE prove to behave as well defined drugs, they may represent the most potent inhibitors of a protein/RNA interaction described, so far.

The importance of LARP6 regulation of type I collagen expression for fibrosis development has been established in several models [20, 22, 25–31]. Here we show that ATO-OA NE can inhibit secretion of type I procollagen by HSCs (cells involved in liver fibrosis, Fig 8) and HLFs (cells involved in pulmonary fibrosis, Fig 9). ATO-OA NE reduced secretion of procollagen from HSCs at concentration of 125 µg/ml or higher, what is 30-40-fold higher concentration than the IC_50_ for LAM binding *in vitro*. The higher effective concentration in cell-based assays may be due to poor cellular permeability of ATO-OA NE, breakdown of NE inside the cells or a weaker inhibitory effect in the cellular environment.

The intracellular retention of type I procollagen was seen at the same concentrations of ATO-OA NE that suppressed the secretion of type I procollagen. Intracellularly, type I procollagen accumulated in large clumps in the perinuclear region (Fig 9B). Normally, type I procollagen is assembled at the membrane of endoplasmic reticulum in well-defined bodies called collagenosomes [38]. This process is LARP6 dependent, so it is likely that ATO-OA NE, by inhibiting LARP6, disrupted the assembly of collagenosomes and caused procollagen clumping into the aggregates that cannot be efficiently exported out of the cells.

The activity of ATO-OA NE in cells is dependent on functional LARP6. In HLFs in which LARP6 had been knocked out there was no effect of ATO-OA NE on intracellular accumulation of type I procollagen (Fig 9C). Likewise, in MEFs where COL1A1 polypeptide is translated from the mRNA without 5’SL and, therefore, not LARP6 regulated [31], there is no effect of ATO-OA NE on type I procollagen biosynthesis (Fig 10). The result in two independent models strongly indicates that the inhibition by ATO-OA NE is due to inactivation of LARP6 and that ATO-OA NE specifically target LARP6 regulated collagen biosynthesis. Constitutive type I collagen expression does not require LARP6 and the assumption is that it will not be affected by ATO-OA NE. It has been reported that LARP6 may bind some other mRNAs [21] or have a general role in translation [19], however, the affinity of binding to these other mRNAs and the molecular recognition involved may be different. Nonetheless, the effect of ATO-OA NE on LARP6 binding to other mRNAs remains to be assessed. LARP6 has a role in cilia formation during embryonic development [63], but this role does not require RNA binding.

Organoids of human pancreatic adenocarcinoma cells express type I collagen [64]. These organoids grow in a 3D matrix, but no effect of ATO-OA NE on the organoids growth rate was seen. In untreated organoids type I procollagen accumulated in intensely stained spots (Fig 11, top panels), but when treated with ATO-OA NE these spots disappeared and only diffuse staining of procollagen was seen (Fig 11, lower panels). We estimated about 3-fold reduction of high intensity procollagen staining with 62.5 µg/ml of ATO-OA NE. This result suggested that ATO-OA NE is effective in suppressing excessive type I procollagen production in human organoids and validated it as a potent scaffold for development of specific antifibrotic drugs.

## Material and methods

### Sources of cefixime

Cefiximine was purchased from different sources; cefixime A from Sequoia Research Products, LTD, cefixime B from Alfa Aesar Chemicals and cefixime C from Research Products International. The drug was dissolved in DMSO at 40 mM and stored in aliquots at -20°C.

### Solvent shift fractionation of cefixime preparations

Cefixime A was dissolved in DMSO at 40 mM and 60 µl of water was added to 100 µl of cefixime solution. The precipitate was collected and redissolved in 100 µl DMSO as fraction 1. Subsequent fractions were obtained by consecutive additions of 10 µl water to the supernatant of the previous fraction and collection of the precipitate, which was dissolved in DMSO.

Additional purification of ATO-OA NE from fractions 2 and 3 was done by adding 19 volumes of methanol to 1 volume of DMSO solution of fractions 2 or 3 and collecting the precipitate. After washing in methanol, the precipitate was dissolved in phosphate buffer pH6. The concentration of ATO-OA NE was determined by weighing in the dry pellets and expressed as µg/ml.

### Mass spectrometry

Cefixime A or the fractions obtained by solvent shift were diluted with methanol, acidified with formic acid and analyzed by Thermo Scientific Q Exactive HF Hybrid Quadrupole-Orbitrap Mass Spectrometer.

### Recombinant proteins and gel mobility shif t experiments

Constructs for expression of recombinant LA, RRM and LAM and their purification have been described [35, 65]. The purity of proteins was estimated by Coomassie staining. For gel mobility shift experiments, 40 nM of fluorescein labeled A1 RNA or Cy5 labeled A2 RNA was incubated with 80 nM of recombinant LAM or LA in 50 µl of 100 mM NaCl, 10 mM Tris pH 7.5, 5 mM MgCl_2_. The reactions were resolved on 6% native acrylamide gels and the gels were imaged by Biorad Imager.

For competition experiments, LA, RRM, BSA or C3 protease were supplemented to the binding reactions at the amounts indicated and after assembling the LAM/5’SL RNA, as above.

### Dynamic light scattering (DLS) measurements

ATO-OA NE sample at 10.4 µg/ml was measured in Wyatt DynaPro Nanostar DLS instrument in duplicate and the hydrodynamic radius was plotted as function of intensity. Conversion from intensity to mass distribution was done by instrument software.

### Fluorescence polarization (FP)

FP was done as described [48]. 80 nM of recombinant LAM or LA was bound to 40 nM of fluorescently labeled A1 5’SL RNA in 100 mM NaCl, 10 mM Tris pH 7.5, 5 mM MgCl_2_ and indicated concentrations of ATO-OA NE or cefixime C were added. 25ul reactions were transferred into 384-well plates and FP was read in Biotek Cytation H1 plate reader. Blank readings with A1 RNA alone were subtracted from the LAM/A1 RNA readings.

### E. coli growth inhibition

E. coli was seeded in 96-well plates at OD_600_ of 0.04 and cefixime or ATO-OA NE were added at the indicated concentrations. Control wells contained no drug. After o/n incubation at 37°C, OD_600_ was measured in 6 replicate wells and shown as average+-1SD.

### Measurement of tryptophan fluorescence

Recombinant LA, LAM or RRM were diluted to 1.4 µM in 400 µl of 100 mM NaCl, 10 mM Tris pH 7.5, 5 mM MgCl_2_ and fluorescence emission spectra were measured after excitation at 288 nM. The spectra were recorded in Cary Eclipse fluorescent spectrometer with proteins alone and after titration with the indicated amounts of ATO-OA NE or cefixime.

### Cell culture and western blots

Human hepatic stellate cells (HSCs), cell line LX-2 [52], human lung fibroblasts (HLFs) [34] and mouse embryonic fibroblasts [31] were grown under standard conditions. ∼50% confluent cells in 12-well plates were treated with the indicated concentrations of ATO-OA NE for 24h or 48h, when the cells were fixed and immunostained. For western blots, after ATO-OA NE treatment the cell medium was replaced by serum free medium, and incubation continued for 3h. After the 3h incubation, serum free medium was collected and an aliquot analyzed by western blot. For western blot, the samples were resolved by 7.5% SDS-PAGE, transferred onto nitrocellulose membrane and probed with anti-COL1A1 antibody (Rockland).

### Immunostaining

The cells in 12-well plates were fixed with 3.7% formaldehyde/0.05% glutaraldehyde/0.5% Triton X-100, blocked with 10% BSA and incubated with 1:10 dilution of anti-collagen antibody (Rockland) overnight. After washing, the Cy3 conjugated secondary antibody at 1:50 dilution was incubated for 1h. Cells were washed, 1 ml PBS was added per well and images were taken at 20x magnification with Cytation 5 imaging reader (Biotek).

For immunostaining of organoids, organoids of human pancreatic adenocarcinoma cells were grown in Matrigel, as described [54]. After two days in culture organoids were treated with 62.5 µg/ml of ATO-OA NE for 3 days, fixed in paraformaldehyde and permeabilized with Triton X-100. Actin was visualized by staining with fluorescent phalloidin and immunostaining for type I collagen was done using anti-COL1A1 antibody (Rockland). Stacks of confocal images of organoids were obtained by Cytation 5 imaging reader, and the maximal projections of stacks were assembled for three viewing fields. Pixel analysis of the maximal projections was done by Image J and pixel count plugin. The threshold for green pixels was set at 30 and the threshold for red pixels was set at 100 and the number of red pixels (procollagen) was normalized to the number of green pixels (actin). Comparison of the normalized pixels was done by paired t-test.

## Acknowledgment

Many thanks to Hyeje Sumajit and Dr. Jerome Irianto for growing and immunostaining of the organoids. Expert technical assistance of Lela Stefanovic is also acknowledged. This study was funded in part by a grant from Live Like Bella Foundation.

